# DUX4-stimulated genes define an antiviral defense program in human placental trophoblasts

**DOI:** 10.1101/2025.01.22.634317

**Authors:** Joshua Hatterschide, Liheng Yang, Carolyn B. Coyne

**Affiliations:** Department of Integrative Immunobiology, Duke University School of Medicine, Durham, NC, USA, Duke University Medical Center, Durham, NC, USA

## Abstract

The placenta combats mother-to-fetus transmission of viruses through the antiviral activities of fetal-derived trophoblasts. Placental trophoblasts employ specialized antiviral strategies to protect against infection while preventing maternal immune rejection of the fetus. However, the full extent of how trophoblasts respond to viral infections is not well understood. To address this, we defined the transcriptional landscape of human trophoblast organoids infected with seven diverse teratogenic viruses. We found that herpesviruses, including HSV-1, HSV-2, and HCMV, did not trigger a typical interferon (IFN) response. Instead, they activated the expression of DUX4 and its downstream target genes, collectively known as DUX4-stimulated genes (DSGs). This program was highly specific for trophoblasts and was associated with cells containing low viral transcripts following HSV-1 infection, suggesting an antiviral activity. Screening of highly expressed DSGs revealed that many of them exhibited anti-herpesvirus activity, indicating the existence of an alternative antiviral pathway similar to the IFN-stimulated gene response. These findings identify DUX4 as a master regulator of a coordinated antiviral program in trophoblasts, specifically targeting a prominent family of teratogenic viruses.

**Graphical Abstract:** 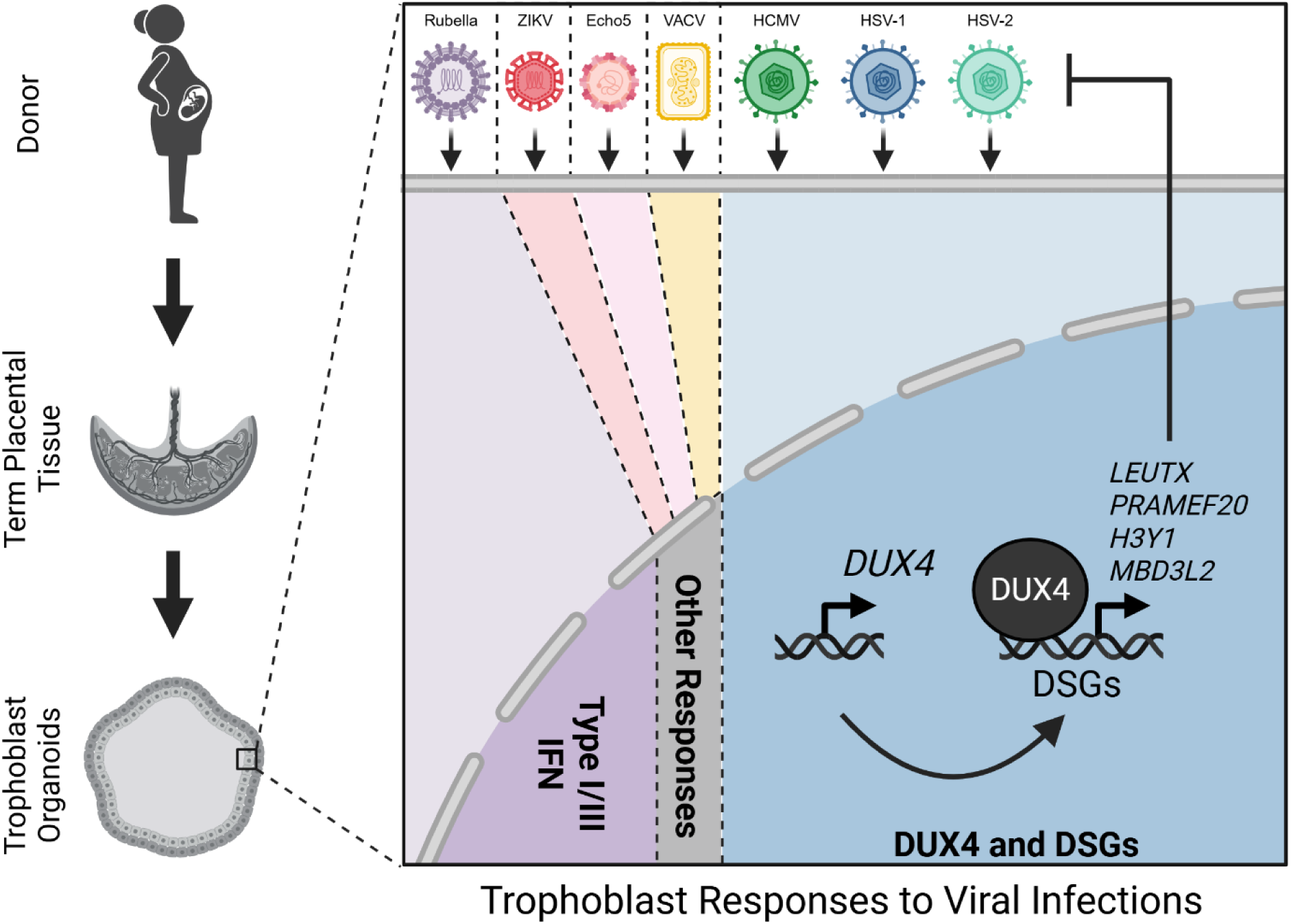

## Introduction

In addition to its critical role in facilitating the exchange of nutrients, gases, and waste between mother and fetus, the placenta acts as a robust barrier against infections, protecting the vulnerable fetus from vertical pathogen transmission during gestation.^1,2^ Certain pathogens, collectively referred to as TORCH pathogens (<u>T</u>oxoplasma, <u>O</u>ther (e.g., enteroviruses, parvovirus B19, hepatitis B virus, syphilis, and Zika virus (ZIKV)), <u>R</u>ubella (RuV), <u>C</u>ytomegalovirus (HCMV), and <u>H</u>erpesviruses) are able to bypass the defenses at the maternal-fetal interface (MFI) and cause placental and/or fetal infections. These infections can lead to a variety of pregnancy complications ranging in severity and including intrauterine growth restriction (IUGR), neurodevelopmental deficits, and loss of the pregnancy.^3–7^ While some TORCH pathogens are vaccine preventable and consequently rare, others remain common. For example, HCMV is estimated to be transmitted from mother to fetus in 1 out of every 200 pregnancies.^8,9^ However, even in the context of a common congenital pathogen such as HCMV, most primary maternal infections during pregnancy are not transmitted to the fetus, demonstrating the efficacy of the immunological defenses present at the MFI.^10^

Fetal-derived trophoblasts form the barrier between maternal and fetal compartments at the MFI, and the antiviral properties of these cells render the placenta resistant to viral infections.^11,12^ Chorionic villi of the human placenta contain several distinct subsets of trophoblasts including cytotrophoblasts (CTBs), extravillous trophoblasts (EVTs), and the syncytiotrophoblast (STB). The multinucleated STB is a single fused cell that covers the entire surface of the placenta and is in direct contact with maternal blood. CTBs are proliferative cells that lie beneath the STB and undergo fusion to grow and repair the STB layer. EVTs arise from columns of CTBs at the tips of chorionic villi, anchor the placenta to the maternal decidua (the pregnancy-specialized maternal endometrium), and facilitate the redirection of maternal blood to the intervillous space.^13^ CTBs, EVTs, and the STB are differentially susceptible to viral infection and thus may respond distinctly to infections.^2^ Despite their critical role as the cellular barrier at the MFI, there remains an incomplete understanding of how human trophoblasts respond to and defend against viral infections.

Responses to pathogens at the MFI must be tightly controlled, as unchecked immune signaling, such as type I interferon (IFN) activation, is linked to adverse pregnancy outcomes.^14–17^ Possibly in response to this selective pressure, trophoblasts have evolved unique antimicrobial properties. For instance, the STB constitutively secretes IFN-λs, which can act in both autocrine and paracrine manners to promote IFN-stimulated gene (ISG) expression to restrict virus replication in the maternal and fetal compartments of the placenta.^12,18–20^ While IFN expression in most cell types is only observed after the detection of pathogen associated molecular patterns (PAMPs), constitutive release of IFNs from trophoblasts is stimulated by viral mimicry by host-derived C19MC Alu dsRNA.^19^ Evidence suggests that trophoblasts also possess IFN-independent mechanisms of virus restriction. Although permissive to HCMV entry, trophoblast organoids (TOs) and trophoblast stem cells have recently been shown to limit HCMV replication at the stage of immediate early gene expression.^20,21^ This restriction occurs even in the presence of ruxolitinib, a JAK inhibitor that inhibits both type I and type III IFN signaling.^20^ However, the role of IFN-independent pathways in virus restriction in trophoblasts remains largely unexplored, highlighting a critical gap in our understanding.

Gaps in our knowledge surrounding how human trophoblasts respond to and defend against virus infection can be attributed to the paucity of tractable systems to model the human MFI. The anatomy of the MFI in many animal models is divergent from that of humans,^22,23^ and animal models also often require the use species-specific viruses (as in the case for HCMV)^24,25^ or immunodeficient or humanized animal models (as in the case of ZIKV in mice).^26^ Human placental explants,^27,28^ isolated primary human trophoblast cells,^12^ choriocarcinoma cell lines,^29^ and immortalized EVT cell lines^30^ have all been employed to study human pathogens in their natural host cells. However, these models either have a limited window of viability or lack physiological properties of *in vivo* trophoblasts.^31^ Trophoblast organoids (TOs) are an *in vitro* model of the placenta that is able to overcome some of the limitations of previous models. TOs and matched maternal decidua gland organoids (DOs) can be derived from chorionic villi at early and late stages of gestation.^20,32,33^ TOs spontaneously differentiate into EVTs and the STB and constitutively secrete hormones such as human chorionic gonadotropin (hCG) and immune molecules such as IFN-λs that are naturally secreted by trophoblasts *in vivo*.^20^

In this study, we profiled the transcriptional responses of TOs to seven teratogenic viruses to define cell-intrinsic mechanisms of virus restriction in trophoblasts and to determine whether these cells responded in a virus-specific manner. We found that TOs mounted distinct responses to RNA and DNA viruses, with the transcription factor DUX4 dominating the response to herpesviruses. DUX4 is a primate-specific pioneering transcription factor that participates in zygotic genome activation and is typically restricted to the 4-8 cells stages of embryonic development.^34^ DUX4 is heterochromatinized in adult somatic cells, and aberrant expression of DUX4 in adult muscle cells is the cause of facioscapulohumeral muscular dystrophy (FSHD).^35,36^ Comparative transcriptional profiling in TOs and DOs derived from matched donor tissue infected with HCMV and HSV-1 revealed that the DUX4 program is enriched in TOs, which occurs independent of constitutive type III IFN signaling. Using single cell RNA sequencing (scRNAseq), we characterized the DUX4 program in trophoblasts in the context of both an inducible DUX4 expression system and HSV-1 infection. DUX4 and accompanying DUX4-stimulated genes (DSGs) were observed to define a low viral gene expression trajectory in HSV-1 infected TOs. To explore the possible antiviral activity of DSGs, we performed high-content screening of the 30 top-most induced DSGs in TOs for antiviral activity against herpesviruses in naturally permissive cells. Analogous to ISGs in the IFN pathway, we observed redundancy in the antiviral activity of DSGs. Expression of DUX4 has been observed following herpesvirus infection in other systems and a few DSGs were previously reported to restrict herpesvirus replication.^37,38^ However, these data shift our understanding of DUX4 and DSGs from a few isolated antiviral genes to a concerted antiviral signaling program that retains key hallmarks of ISGs, including a robust and overlapping antiviral gene induction mechanism. Together, these findings unveil a human trophoblast-enriched anti-herpesvirus pathway that mirrors the functional principles of the IFN response, providing new insights into host defense strategies against viral infections at the maternal-fetal interface.

## Results

### Human trophoblast organoids respond distinctly to diverse viruses

To directly compare how human placental trophoblasts respond to infection with teratogenic viruses, TOs from two male donors and one female donor were infected with seven teratogenic viruses that belong to diverse virus families. These viruses included three herpesviruses: HCMV, HSV-1, HSV-2, a poxvirus: Vaccinia virus (VACV), and three positive sense RNA viruses: echovirus 5 (Echo5), rubella virus (RuV), and Zika virus (ZIKV). RNA was collected from mock and infected TOs and sent for polyA-selection bulk RNA sequencing. Reads were mapped to both the human and viral genomes. We observed minimal mapping of reads to the viral genomes in mock samples and little donor-to-donor variation in viral read levels in infected samples **(Figure S1A)**. We have previously reported that HCMV replication is impaired in TOs. However, entry and gene expression were confirmed based on high quantities of reads mapping to the HCMV genome **(Figure S1A)**. In contrast, several of the other viruses exhibited evidence of replication, albeit at low levels in some cases. Fluorescent protein expression from HSV-1 (GFP) was detectable by fluorescence microscopy, and HSV-1 and HSV-2 both induced cytopathic effect (CPE) in TOs **(Figure S1B)**. Fluorescent tag expression from VACV (YFP) was also visible by fluorescence microscopy **(Figure S1C)**. CPE was observed in TOs following RuV infection, but not following infection with either Echo5 or ZIKV **(Figure S1C-D)**. Consistent with this observation, RuV was the only positive sense RNA viruses that yielded high levels of both positive and negative strand genomic reads in all donors, suggesting it is the only one of these viruses readily engaging in productive replication **(Figure S1E)**. For the remaining analyses, reads mapping to the virus genomes were excluded to focus on host cell responses.

Differential gene expression analysis was conducted using the DESeq2 package, incorporating donor as a factor in the statistical model to account for variability between individual donors.^39^ Differentially expressed genes (DEGs) were defined as host genes with Log_2_ (fold-change) > 1 and adjusted *p*-value < 0.05. The differences in magnitude of the induced responses ranged from zero DEGs following ZIKV infection to 5453 DEGs following VACV infection **(Figure 1A-G and Table S1)**. We also observed variability in the directionality of responses. HSV-1, HSV-2, and RuV all led to substantially more upregulated DEGs than downregulated. HCMV and VACV both led to a relatively even distribution between upregulated and downregulated DEGs. Echo5 was the only virus with a response that skewed toward downregulation **(Figure 1A-G)**. The small transcriptional responses elicited by Echo5 and ZIKV may arise from impaired virus replication in TOs, which would lead to lower levels of pathogen associated molecular pattern (PAMP) detection and a diminished transcriptional response. This is consistent with the observation that these viruses had the lowest quantities of viral reads **(Figure S1A and E)**. Pairwise comparisons between each infection condition revealed some overlap in upregulated and downregulated DEGs between these diverse viruses **(Figure 1H).** However, after excluding Echo5 and ZIKV (due to their limited responses), only two DEGs were upregulated by all of the five remaining viruses: *EGR3* and *THBS1*. This suggests that TOs respond distinctly to infection in a virus-specific manner.

**Figure 1.**
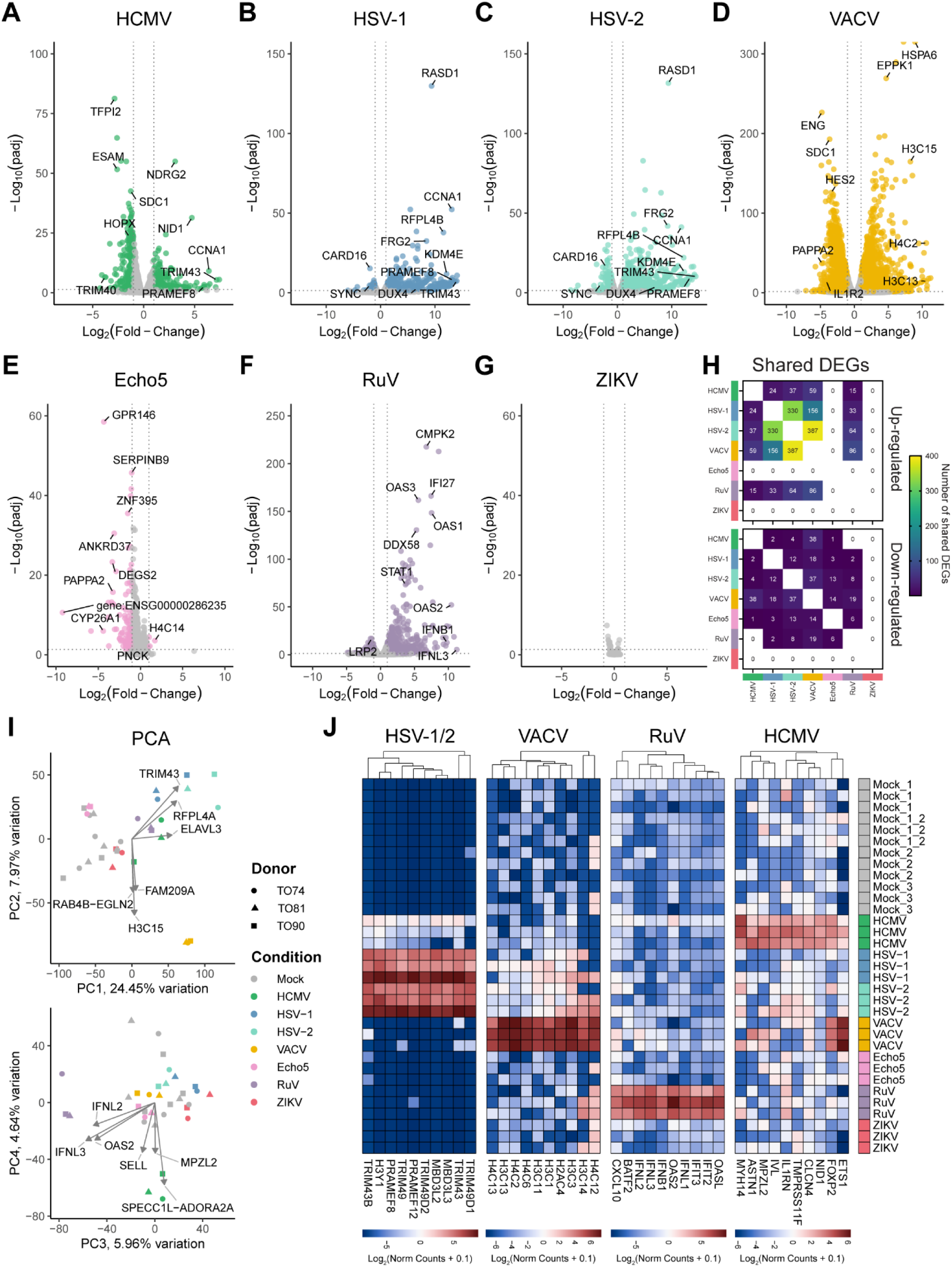
Trophoblast organoids respond distinctly to diverse viruses. Trophoblast organoids from three donors were infected with HCMV, HSV-1, HSV-2, VACV, Echo5, RuV, and ZIKV at an MOI of 2. VACV and Echo5 infections were performed in suspension on organoids released from matrigel that were re-imbedded in matrigel following inoculation. All other infections were performed on spherical organoids grown on matrigel coats. mRNA was purified and analyzed by bulk RNAseq. **(A-G)** Volcano plots portraying the DEGs relative to mock samples. Colored circles represent significant DEGs (Log2(fold-change) > 1 and adjusted *p*-value < 0.05). Gray circles represent genes whose expression was not significantly impacted. Also see **Table S1**. **(H)** Heatmap showing the number of shared DEGs between each infection condition. Genes that were found to be differentially expressed between donors, timepoints, or experimental sets in the mock infected samples were excluded from these tallies. **(I)** PCA of RNAseq data from each sample is portrayed in two graphs: principle component 1 (PC1) vs PC2 (top) and PC3 vs PC4 (bottom). Shapes denote the donor a sample came from, and color denotes the infection condition of the sample. **(J)** Heatmaps portraying the genes that define the responses unique to virus types. Values are log_2_ normalized counts from DESeq2 analysis plus a pseudo count of 0.1. Keys are at the bottom of each chart. The scale is not constant between each heatmap. Red indicates higher expression and blue indicates lower expression. Hierarchical clustering of genes is shown at the top.

To further categorize these responses, we performed a principal component analysis (PCA). Four groups were found to segregate from the mock-infected samples along the first four principle components: HSV-1- and HSV-2-infected, VACV-infected, RuV-infected, and HCMV-infected **(Figure 1I)**. As expected, Echo5- and ZIKV-infected samples clustered closely with the mock-infected samples. We next analyzed the loadings from the PCA to identify sets of genes that defined these unique responses. The response to VACV was predominantly defined by genes encoding histone variants such as *H3C1* and *H4C2* **(Figure 1J, middle left)**. RuV was the only virus that induced type-I and type-III IFN genes such as *IFNB1* and *IFNL1* along with downstream IFN-stimulated genes (ISGs) including *OASL*, *IFIT2,* and many others **(Figure 1J, middle right)**. The unique response to HCMV was defined by genes including *IVL*, *IL1RN*, and *MPZL2* **(Figure 1J, right)**. HSV-1- and HSV-2-infected samples were defined by tripartite motif family genes such as *TRIM43* and *TRIM49*, and preferentially expressed antigen of melanoma family genes such as *PRAMEF8* and *PRAMEF12* **(Figure 1J, left)**.

Many of the genes that defined the HSV-1- and HSV-2-infected samples were also induced by HCMV. In fact, there were 28 genes induced by all three herpesviruses **(Figure S1B)**. Notably, most of these shared DEGs were induced by herpesvirus infections and were not induced by any other viruses **(Figure S1C)**. These findings suggest the existence of a herpesvirus-specific pathway activated in trophoblasts.

### Trophoblasts respond to herpesviruses with DUX4-stimulated genes

Several of the shared differentially expressed genes induced by HSV-1, HSV-2 and HCMV, such as *TRIM43*, are putative targets of the transcription factor DUX4. DUX4 is a pioneering transcription factor that is normally expressed during embryogenesis between the four-cell and eight-cell stages^34^ and is pathogenic when aberrantly expressed in adult muscle cells.^35,36^ DUX4 has also been shown to be upregulated following HSV-1 infection.^37^ In the TO infection bulk RNAseq analyses, *DUX4* expression was up-regulated in all donors by HSV-1 and HSV-2 and by two donors by HCMV **(Figure 2A)**. DUX4 was not detected in any other infection conditions. However, *DUX4* expression is notoriously difficult to detect even using next generation sequencing methods.^40,41^ Therefore, we confirmed the expression of *DUX4* and the DUX4 transcriptional-target, *TRIM43*, only in herpesvirus-infected samples in all three donors using qRT-PCR **(Figure S2A-B)**.

**Figure 2.**
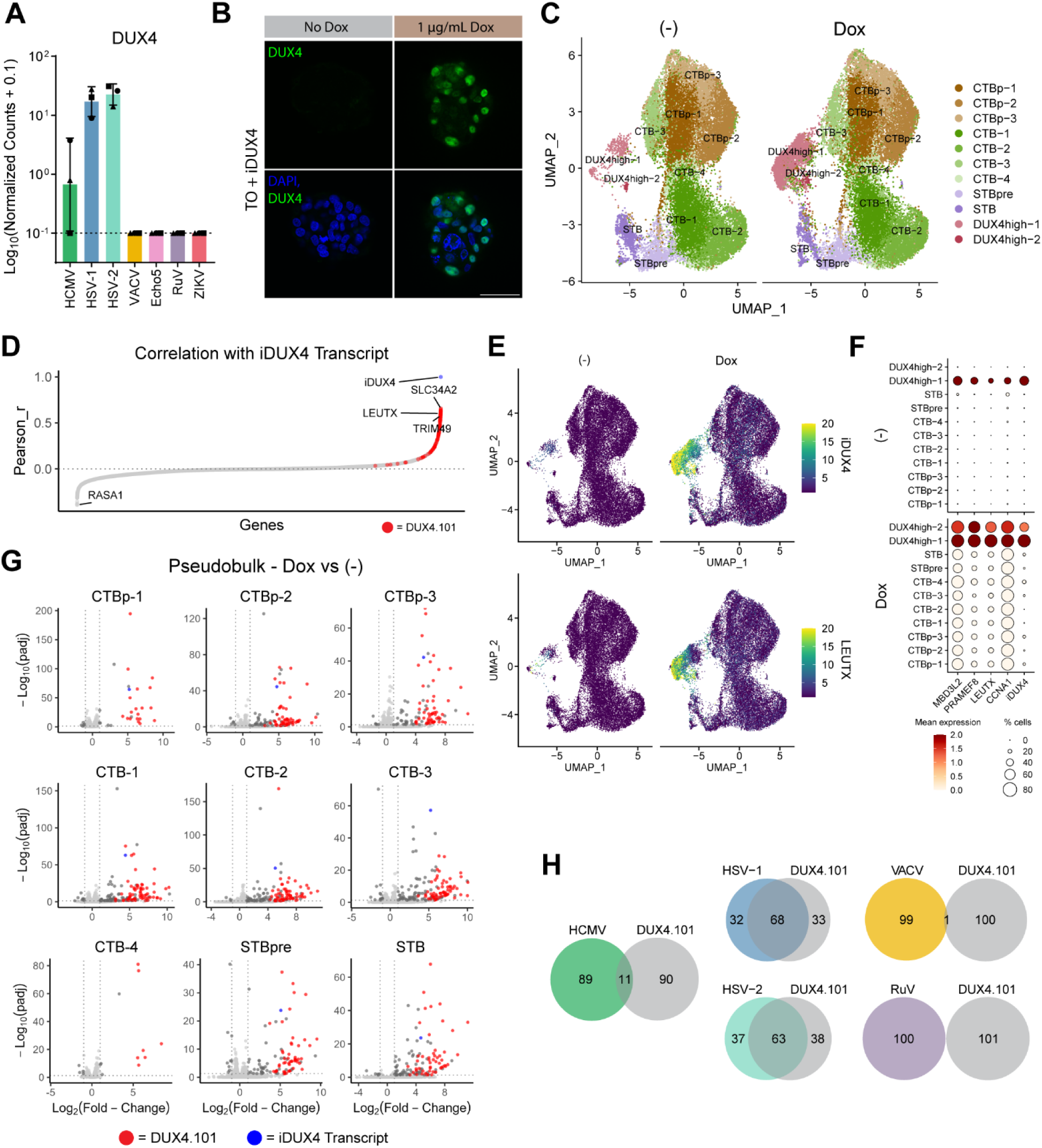
Trophoblasts respond to herpesviruses with DUX4-stimulated genes. **(A)** Bar plot portraying the expression of *DUX4* in the RNAseq experiments from Figure 1. Shapes denote the donor of each data point. **(B)** Confocal micrographs of TOs engineered to express DUX4 under a doxycycline (Dox) inducible promoter. DUX4 is displayed in green and DAPI-stained nuclei are in blue. Scale bar = 100 μm. **(C)** UMAP of clusters from three donors of mock-treated TOs and three donors of Dox-treated TOs. Clusters are labeled as proliferative cytotrophoblasts (CTBp), cytotrophoblasts (CTB), pre-syncytiotrophoblasts (STBpre), syncytiotrophoblasts (STB), or DUX4-high cells. Multiple clusters from a single cell type are annotate with a -number after the cluster name. **(D)** Plot portraying the correlation (Pearson’s r) between iDUX4 transcript and all other measured genes. Values are plotted in rank order. Genes that belong to the DUX4.101 list are displayed as red circles. Also see **Table S2**. **(E)** Feature plots of iDUX4 (top) and the *LEUTX* (bottom) expression. Keys are shown to the right. **(F)** Dot plots showing the average expression (ALRA assay) and proportion of cells expressing selected DSGs and iDUX4 in each cluster separated into mock-treated samples (top) and Dox-treated samples (bottom). **(G)** Pseudobulk analysis identifying significant DEGs (Log2(fold-change) > 1 and adjusted *p*-value < 0.05) between Dox-treated samples and mock-treated samples. Results from all clusters other than the DUX4-high clusters are shown. Genes that belong to the DUX4.101 list are displayed as red circles and iDUX4 transcript is displayed as a blue circle. Also see **Table S4**. **(H)** Venn diagrams denoting the overlap between the top 100 DEGs upregulated by each virus and the DUX4.101 list.

DUX4 has never been studied in trophoblasts, and DSGs in these cells have yet to be defined. To characterize DSGs in trophoblasts, we developed a model of DUX4 expression in TOs. TOs from three donors were transduced with a lentiviral vector encoding codon-optimized DUX4 under a doxycycline-inducible promoter (iDUX4). Transduced organoids were selected with puromycin, and expression of DUX4 upon treatment with doxycycline was confirmed by immunofluorescence **(Figure 2B)**. One strength of the TO model is that they retain the capacity to differentiated into several lineages of trophoblasts present in the placenta *in vivo*. To define DSGs across trophoblast cell types, and to determine whether there were trophoblast cell type-specific DSGs, doxycycline-treated and mock-treated organoids from three donors were dissociated and analyzed by single-cell (sc)-RNAseq. A total of 23,981 doxycycline-treated cells and 23,233 mock-treated cells passed quality control and were used for further transcriptional analysis following integration to correct for donor and batch effects. Cluster analysis revealed 11 cell clusters that were present in both conditions **(Figure 2C and Figure S2C)**. The identities of the clusters were defined by the expression of canonical trophoblast marker genes^42–44^ **(Figure S2E-F)**. The proliferative CTB clusters (CTBp-1, -2, and -3) were defined by high expression of *MKI67*, *CCNA2*, *PCNA*, and *TOP2A*. CTB clusters (CTB-1, -2, -3, and -4) were defined by cells that express high levels of the pan-trophoblast markers *PAGE4*, *PEG10*, *SMAGP*, and *TP63*, but lacked expression of more specialized programs. Of note, CTBp-1 and CTB-4 also expressed markers of cell column CTBs such as *ITGB6*, *ATF3*, *PLAC8*, and *PLAU*. Cells in the process of fusing into the STB (STBpre) were defined by the expression of the endogenous retroviral elements that mediate and modulate fusion such as *ERVV-1*, *ERVFRD-1*, *ERVW-1*, and *ERVE-1*. Markers of STB function including *HOPX*, *SDC1*, *CGB5*, and *PSG3* were used to define the STB cluster. Two clusters were classified based on high iDUX4 expression in the doxycycline-treated samples (DUX4-high-1 and DUX4-high-2). Based on the expression of other markers, these clusters most closely resembled CTBs **(Figure S2E)**. The presence of select cells in the “DUX4-high” clusters in mock-treated samples is likely due to leaky expression of iDUX4. The DUX4-high-1 cluster expands from an average of 1.6% of the mock-treated cells to an average of 11.6% of the doxycycline-treated cells **(Figure S2D)**. The remaining clusters occupied a similar proportion of cells between donors and conditions.

We first sought to identify genes that were broadly controlled by DUX4 across cells in the dataset. We calculated the Pearson’s correlation coefficient (r) between the expression of iDUX4 and other genes across all cells. One hundred and fifty genes were found to positively correlate with iDUX4 expression with r > 0.4 suggesting they were stimulated by iDUX4 and could be classified as DSGs **(Figure 2D and Table S2)**. To test whether these genes were similar to DSGs in other cell types, we generated a list of 101 cell-type-independent DSGs (DUX4.101) by reanalyzing and comparing bulk RNAseq datasets in which DUX4 was overexpressed in iPSCs (a model of DUX4 in embryogenesis)^34^ and MB135 cells (a model of DUX4 in muscular dystrophy)^45^ **(Figure S3A-C and Table S3)**. The expression of many of the genes in the DUX4.101 list strongly correlated with iDUX4, suggesting the DUX4 program in TOs is similar to that of other cell types **(Figure 2D)**. We also confirmed overlap between the DUX4.101 list and genes controlled by DUX4 in TOs by performing bulk RNAseq on doxycycline-treated and mock-treated organoids **(Figure S3D-E and Table S3)**. While there were many genes uniquely modulated by iDUX4 in TOs, these data demonstrate that the canonical DUX4 program is induced in trophoblasts. 98 of the genes that correlated with iDUX4 expression with r > 0.4 were not a part of the iDUX4 list, including *GTF2F1*, *PTP4A1*, *DBR1*, *ODC1*, *SOX15* amongst others.

We next investigated which trophoblast cell types predominantly expressed the DUX4 pathway and whether it could be activated in a trophoblast cell type-specific manner. Expression of genes in the DUX4.101 list were elevated in the doxycycline-treated samples and predominantly found in the two DUX4-high clusters **(Figure 2E-F)**. This demonstrates that cells that express iDUX4 are transcriptionally distinct enough to cluster separately from other trophoblasts. As iDUX4 expression could also be observed in the non-DUX4-high clusters **(Figure 2F)**, we next defined the impact of iDUX4 expression in these cell types. We performed pseudobulk analysis to identify DEGs between the doxycycline-treated and mock-treated samples for each cluster. Although there were cluster-specific differences in DEGs, genes in the DUX4.101 list were found to be among the most highly differentially expressed genes across all clusters **(Figure 2G and Table S4)**. These data demonstrate that many of the canonical DSGs are also induced by DUX4 expression in trophoblasts and that DUX4 expression is capable of driving the expression of these genes in all trophoblast lineages.

Finally, we determined whether any of the DEGs induced in TOs in response to herpesvirus infections were DSGs. We found that the DUX4.101 list significantly overlapped with the top 100 DEGs induced by HCMV, HSV-1, and HSV-2, while overlapping by only one gene with the top DEGs induced by VACV and zero with RuV **(Figure 2H and Figure S3F-K)**. HCMV may have induced fewer DSGs than HSV-1 or HSV-2 due to its impaired replication in trophoblasts ^20,21^. Based on these data, we conclude that many of the shared genes induced by herpesviruses in TOs are DSGs.

### DUX4-stimulated genes are induced to higher levels in trophoblast organoids than maternal-derived decidua epithelial organoids

Trophoblasts restrict HCMV replication.^20,21^ For example, we have previously reported that fluorescently-tagged HCMV strains express high levels of fluorescent protein in maternal DOs while exhibiting minimal tag expression in matched TOs.^20^ Unlike HCMV, HSV-1 exhibits signs of replication in TOs, including fluorescent tag expression and promoting CPE **(Figures S1B)**. To test whether HSV-1 replication is impaired in TOs compared to DOs, both types of organoids were mock-treated or infected with HSV-1 (MOI of 0.1) enabling multiple rounds of replication. Organoid media was sampled at 1, 2, and 3 dpi, and HSV-1 infectious virus was quantified. While TOs indeed supported the production of infectious HSV-1 particles as predicted, we observed an average of 8.4-fold lower titers produced from TOs than matched maternal DOs across time points **(Figure 3A-B)**. These data suggest that TOs restrict HSV-1 replication.

**Figure 3.**
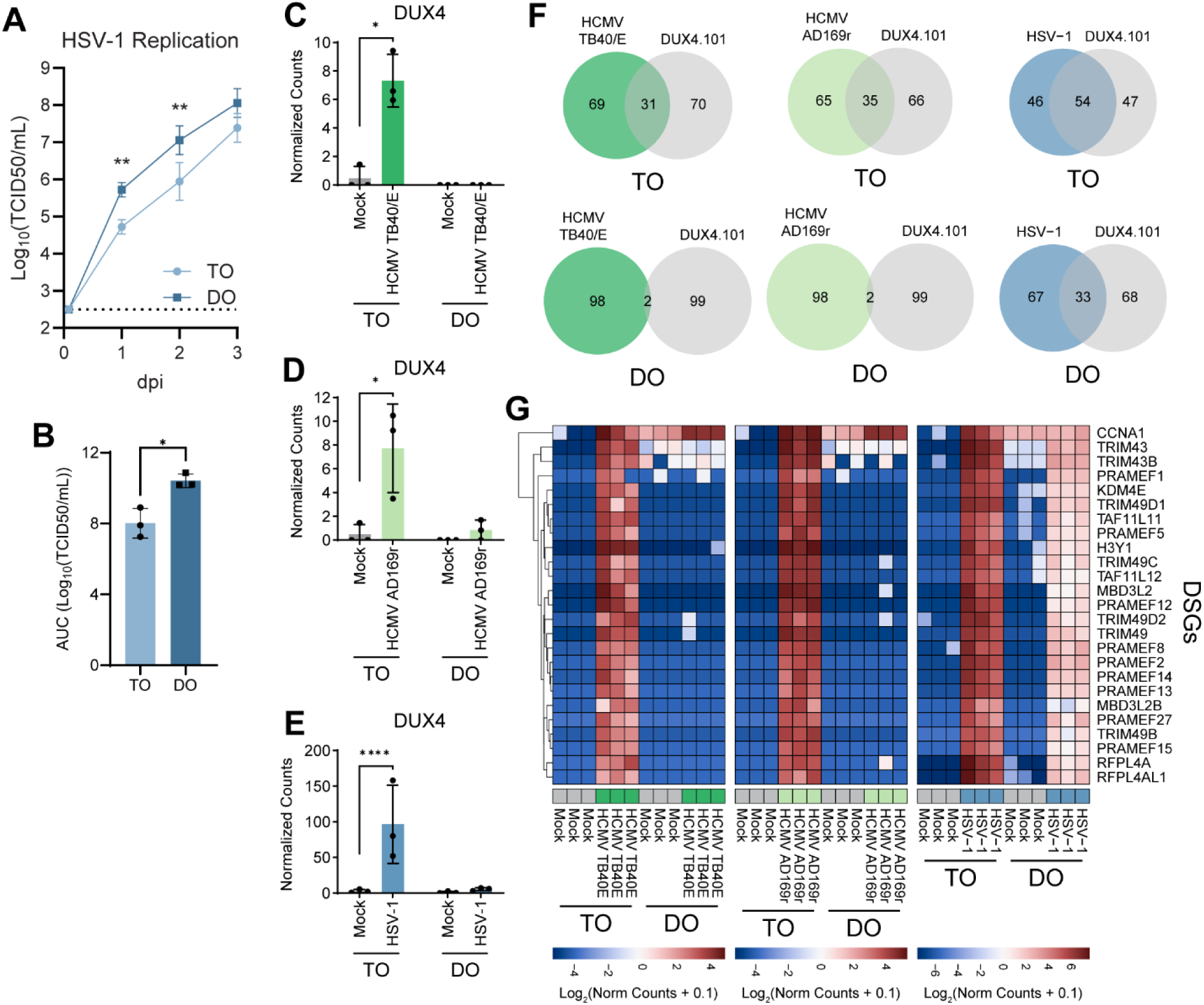
DUX4-stimulated genes are induced to higher levels in trophoblast organoids than maternal-derived decidua epithelial organoids. **(A-B)** TOs and DOs were seeded on matrigel coats and infected with HSV-1 at an MOI of 0.1. Infectious HSV-1 particles were quantified using the 50% tissue culture infectious dose (TCID50) method at 2 hpi as well as 1, 2, and 3 dpi (A). Graph displays the mean ± SD at each timepoint. Statistical significance was calculated using 2-way ANOVA, independent t-tests for each timepoint, and the Šidák correction for multiple comparisons. Area under the curve (AUC) was calculated for each independent series of samples (B). Each individual data point is displayed. Bar graph displays the mean ± SD. Statistical significance was determined using by t-test. **(C-G)** HSV-1 infections were performed in suspension on organoids released from matrigel and subsequently re-imbedded in matrigel. RNAseq analysis was employed to assess if TOs and DOs differentially express DUX4 or DSGs when infected with HCMV TB40/E, HCMV AD169r, and HSV-1 (MOI of 5). Also see **Table S5**. **(C-E)** Bar plots portraying the expression of *DUX4* in TOs and DOs in mock-treated samples or those infected with HCMV TB40/E (C), HCMV AD169r (D), and HSV-1 (E). Each individual normalized count value is displayed. Bars represent mean ± SD. Statistical significance was determined using DESeq2 differential expression analysis. **(F)** Venn diagrams denoting the overlap between the top 100 DEGs upregulated by each virus and the DUX4.101 list with TO infections on the top and DO infections on the bottom. **(G)** Heatmaps portraying the expression of canonical DSGs following infection of TOs and DOs with HCMV TB40/E, HCMV AD169r, and HSV-1. Values are log_2_ normalized counts from DESeq2 analysis plus a pseudo count of 0.1. Keys are at the bottom of each chart. The scale is not constant between each heatmap. Red indicates higher expression and blue indicates lower expression. Hierarchical clustering of genes is shown on the left. * p < 0.05, ** p < 0.1, **** p < 0.0001.

TOs differentially respond to HCMV when compared to maternal DOs, suggesting that there may be trophoblasts-specific responses to herpesvirus infection.^20^ We next assessed whether DUX4 and DSGs were differentially induced in TOs and DOs following herpesvirus infections. Matched TOs and DOs were mock-treated or infected with HSV-1 (MOI of 5). RNA was collected and sent for polyA selection bulk RNA sequencing. Additionally, we reanalyzed published RNA sequencing data from infection of TOs and DOs with the HCMV strains TB40/E and AD169r (a clone of AD169 engineered to express the HCMV pentameric complex).^20,46^ All RNA sequencing datasets were remapped to the human transcriptome following the same pipeline and analyzed using DESeq2 **(Table S5)**. We found that DUX4 expression was more than 9-fold higher in TOs than DOs following infection with HCMV TB40/E **(Figure 3C)**, HCMV AD169r **(Figure 3D)**, and HSV-1 **(Figure 3E)**. Analysis of the top 100 upregulated DEGs following each infection revealed greater overlap between the DEGs of TOs and the DUX4.101 list compared to the DEGs of DOs **(Figure 3F)**. While these DSGs were highly induced in TOs, nearly all of them were not induced in DOs following infection with HCMV TB40/E or HCMV AD169r **(Figure 3G and Figure S4A-D)**. Although DSGs were induced following HSV-1 infection in both TOs and DOs, all were induced between 6- and 50-fold more in TOs **(Figure 3G and Figure S4E-F)**. We used qRT-PCR to confirm enhanced expression of *DUX4* and the DSGs *TRIM43B* and *PRAMEF1* in three donors of TOs compared to three donors of DOs following HCMV TB40/E infection **(Figure S4G-I)**. These data demonstrate that trophoblasts respond with higher expression of *DUX4* and DSGs than maternal decidua epithelial cells.

### Expression of DUX4 and DSGs is not dependent on constitutive type-III IFN signaling

The mechanisms by which DUX4 and DSG expression are induced by herpesviruses remain unknown. Additionally, the signaling pathways that modulate this program to drive its heightened expression in TOs have yet to be identified. Trophoblast stem cells, another *in vitro* model of trophoblasts, can be induced to differentiate into EVTs and the STB under specific culture conditions.^47^ Upon reanalysis of previously published RNA sequencing data from trophoblast stem cell responses to infection with mCherry-tagged HCMV,^21^ we found that infection of STB-differentiated trophoblasts elicited between 4- and 960-fold higher levels of DUX4 and DSGs than mCherry-positive undifferentiated trophoblasts stem cells **(Figure S4J)**. EVT-differentiated trophoblasts infected with HCMV also elicited a greater DUX4 and DSG response than undifferentiated TSCs, but less so than the STB-differentiated condition. These data suggest that trophoblast differentiation, especially towards the STB, may drive the high levels of this program observed in TOs.

Trophoblasts constitutively secrete type III IFNs, which is a major innate immunological difference between TOs and DOs.^20^ Additionally, type III IFNs are produced by the STB.^18^ Although DUX4 and DSG expression has not been previously linked to type-III IFN signaling, it could explain the increased expression of DUX4 and DSGs in TOs over DOs and in the STB over other trophoblasts. To test this, we generated *IFNLR1* knockout (KO) TOs using CRISPR-Cas9 and a single guide RNA (sgRNA) targeting exon three of the *IRNLR1* locus. To validate KO of *IFNLR1* genetically, we amplified the sgRNA target site using high-fidelity PCR and sequenced the product using Sanger sequencing. Using the ICE tool (Synthego) to predict the mixture of insertions and deletions present in the mixture, we found -19 bp and -10 bp deletions >95% of the sequences in the IFNLR1 KO clone **(Figure 4A-B)**. To test whether IFNLR1 KO TOs are responsive to type-III IFNs, parental and IFNLR1 KO organoids were treated with recombinant IFNλ3. RNA was collected and qRT-PCR was used to measure the expression of several ISGs. Whereas *OASL* and *MX1* were induced ∼ 5-fold and 10-fold respectively in the parental TOs treated with IFNλ3, neither was induced in IFNLR1 KO TOs **(Figure 4C-D)**. Additionally, *OASL* and *MX1* were expressed at a significantly lower baseline-level in IFNLR1 KO cells, likely due to disruption of constitutive autocrine type III IFN signaling. IFNLR1 KO and parental cells were infected with HSV-1 to induce the expression of DUX4 and DSGs. RNA was collected from mock and infected organoids and qRT-PCR was used to measure the expression of the HSV-1 immediate early transcript ICP4, *DUX4*, and the DSGs: *TRIM43* and *PRAMEF1*. HSV-1 ICP4 was expressed at similar levels in parental and IFNLR1 KO cells, suggesting that type III IFN signaling does not restrict HSV-1 replication **(Figure 4E)**. *DUX4* expression was increased in IFNLR1 KO TOs and *TRIM43* and *PRAMEF1* were expressed at similar levels in control and KO organoids **(Figure F-H)**. These data suggest that the increased expression of *DUX4* and DSGs in TOs occurs through a mechanism independent of type-III IFN signaling.

**Figure 4.**
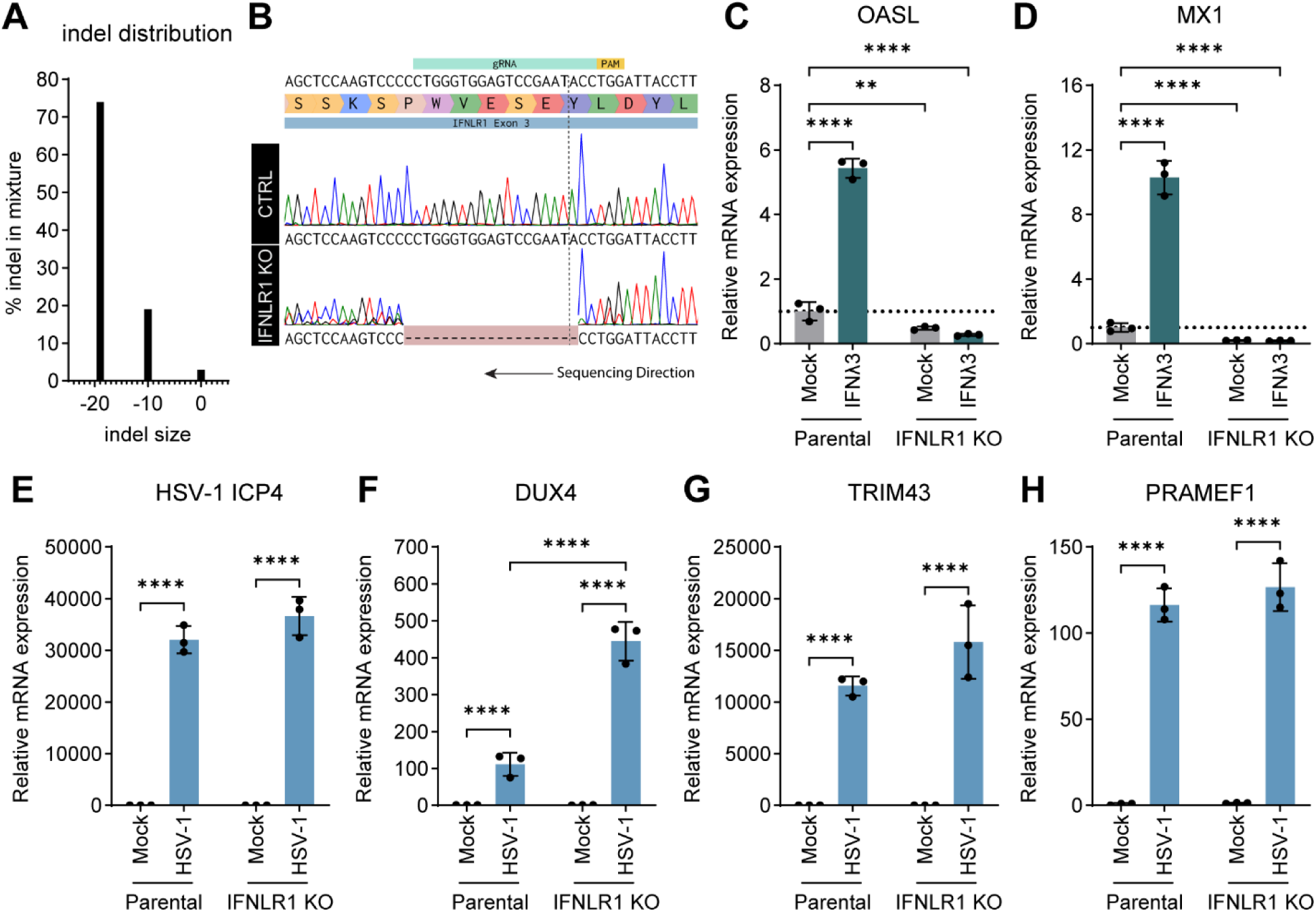
Expression of DUX4 and DSGs are not dependent on constitutive type-III IFN signaling. TOs were transduced with lentiviral vectors engineered to express Cas9 and an sgRNA to target exon three of the human IFNLR1 locus. Single cell cloning was used to isolate homogenous organoid lines. PCR amplicons of the sgRNA target site were sent for sanger sequencing and analyzed using the Synthego ICE tool. **(A)** Predicted indel distribution of the IFNLR1 KO clone. **(B)** Sanger sequencing traces of the IFNLR1 KO clone aligned to a control sequencing trace. Sequencing proceeded from right to left as the traces are displayed. **(C-D)** IFNLR1 KO and parental organoids grown on matrigel coats were mock-treated or treated with 5 ng/μL IFNλ3. qRT-PCR was used to measure the expression of the ISGs *OASL* and *MX1*. **(E-H)** IFNLR1 KO and parental organoids grown on matrigel coats were infected with HSV-1 to induce DUX4/DSGs. qRT-PCR was used to measure the expression of HSV-1 ICP4, *DUX4*, *TRIM43B*, and *PRAMEF1*. Values are plotted as expression relative to parental mock and each individual data point is displayed. Bars display mean ± SD. Statistical significance was determined by ANOVA with the Tukey multiple comparisons tests. * p < 0.05, ** p < 0.01, **** p < 0.0001.

### DUX4 expression is inversely correlated with HSV-1 gene expression in trophoblast organoids

In comparison to matched maternal DOs, TOs restricted the replication of HCMV and HSV-1 and simultaneously expressed substantially higher levels of DUX4 and DSGs **(Figure 3)**. This correlation suggests that these genes may contribute to making TOs less permissive to herpesvirus replication. To explore this hypothesis, we first asked how DUX4 expression in trophoblasts correlated with HSV-1 gene expression. TOs were mock-infected or infected with HSV-1 (MOI 5; GFP tagged to UL25) and then DUX4 expression was imaged by immunofluorescence confocal microscopy. DUX4 protein expression was only observed in infected trophoblasts within TOs, confirming our earlier transcriptomics results **(Figure 5A)**. Additionally, we noted that DUX4 expression was observed primarily in cells with little to no expression of the HSV-1 GFP tag. To confirm this observation, we quantified the nuclear intensity of DUX4 and the HSV-1 GFP tag across thousands of cells and found that DUX4 intensity inversely correlated with HSV-1 GFP intensity **(Figure 5B)**, which was consistent across independent infections **(Figure 5C)**. These data are consistent with the hypothesis that DUX4 and DSGs may contribute to restricting herpesvirus replication in TOs.

**Figure 5.**
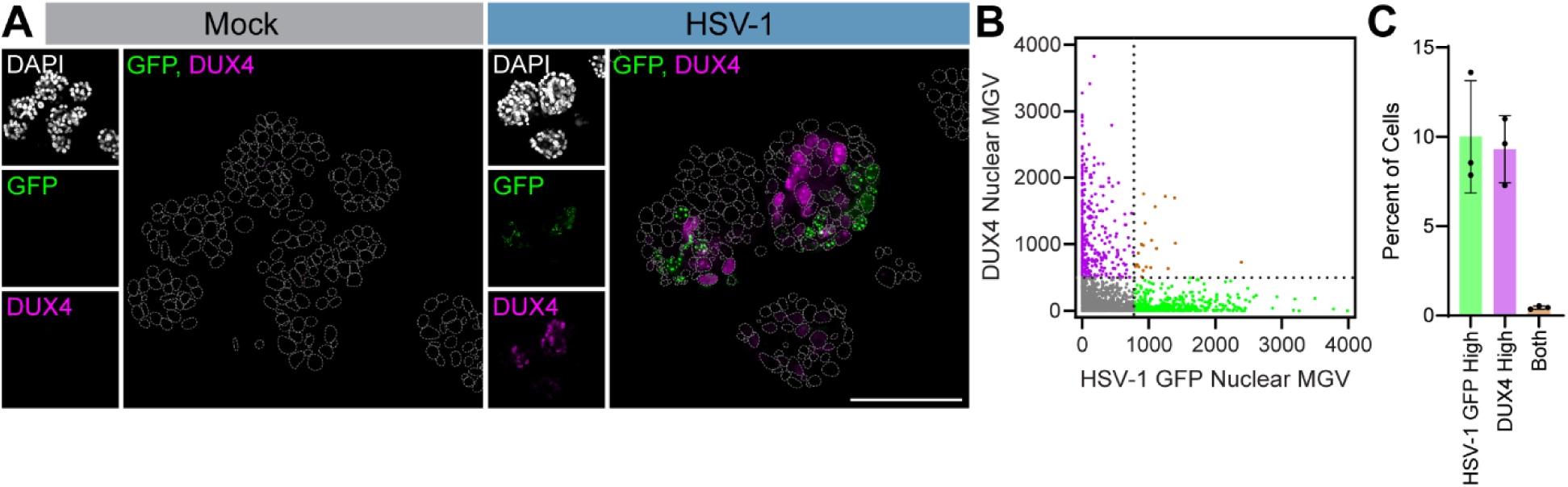
DUX4 expression inversely correlates with HSV-1 gene expression in trophoblast organoids. TOs were grown on coats and infected with HSV-1 (MOI of 5). DUX4 was stained by immunofluorescence and DUX4 and HSV-1 GFP-UL25 were imaged by confocal microscopy. **(A)** Confocal micrographs of mock-infected and HSV-1-infected trophoblast organoids. Individual channels are displayed to the left with DAPI in gray, GFP-UL25 in green, and DUX4 in magenta. Nuclear outlines are shown in the merged image as dashed gray lines. Scale bar = 100 μm. **(B)** Quantification of nuclear mean gray value (MGV) of DUX4 and HSV-1 GFP-UL25. Each dot represents an individual nucleus. **(C)** Bar graph of the proportion of cells determined HSV-1 GFP-high, DUX4-high, or both. Black circles represent each independent datapoint. Bars display mean ± SD.

### DUX4-stimulated genes define a low viral gene expression trajectory in HSV-1-infected trophoblast organoids

To further define the relationship between the DUX4-DSG program and herpesvirus infections in trophoblasts, we next investigated which trophoblast cell types expressed this program in response to infection and examined how the expression of these genes compared to that of viral genes. HSV-1-infected and mock-infected organoids from two donors were dissociated and analyzed by scRNAseq. A total of 14,846 HSV-1-infected cells and 13,881 mock-infected cells passed quality control and were used for further analysis following integration to correct for donor and batch effects. Clustering analysis revealed 11 clusters of cells that were present in both conditions **(Figure 6A and Figure S5A-C)**. The identities of the clusters were defined by the expression of the canonical trophoblast marker genes described above **(Figure S5D-J)**.

**Figure 6.**
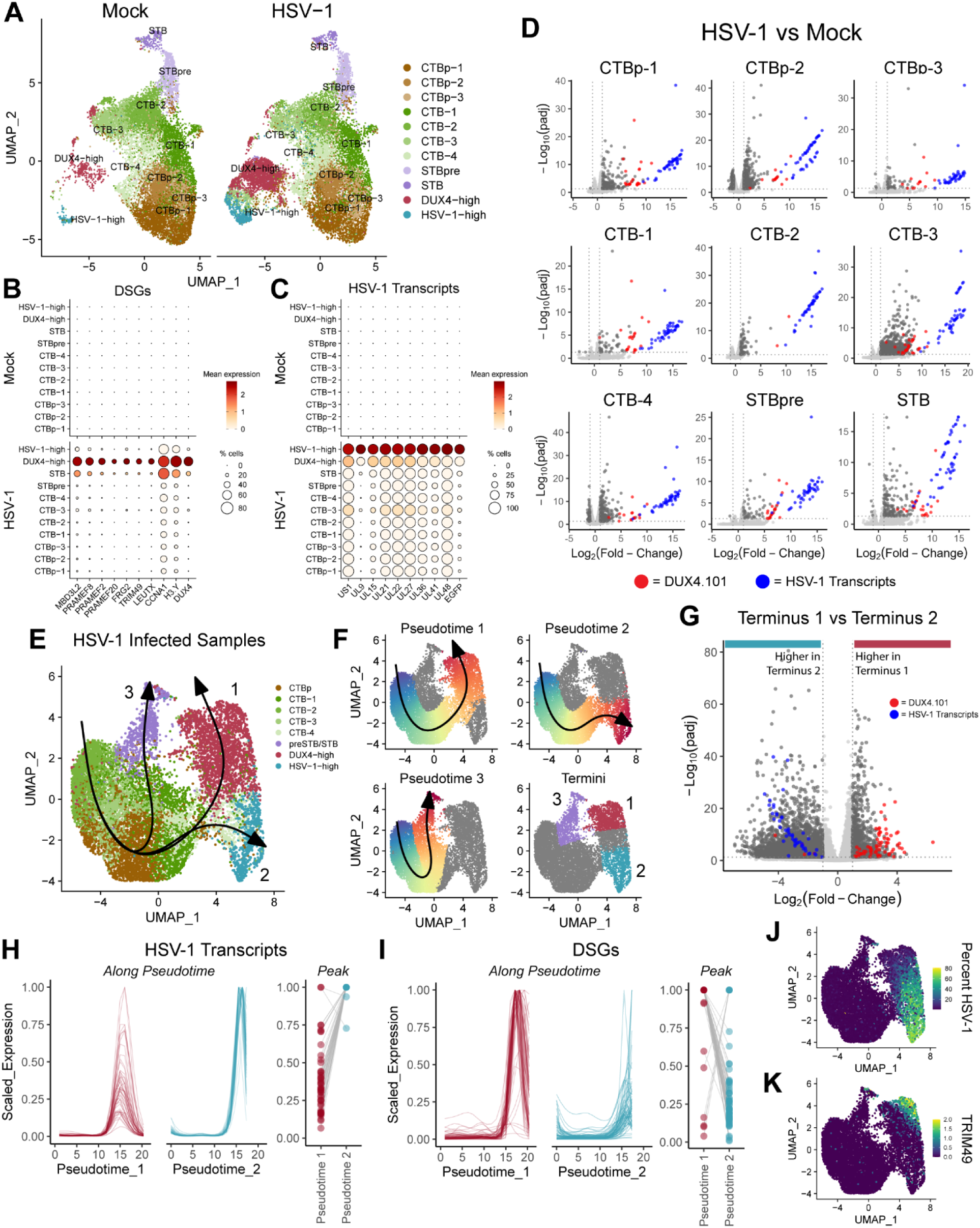
DUX4-stimulated genes define a low viral gene expression trajectory in HSV-1-infected trophoblast organoids. TOs released from matrigel were infected with HSV-1 in suspension then returned to matrigel imbedding. HSV-1-infected and mock-infected organoids from two donors (2x TO74 samples and 1x TO84 sample each) were analyzed by scRNAseq. **(A)** UMAP of clusters from all mock-infected and all HSV-1-infected TOs separated by infection status. Clusters are labeled as proliferative cytotrophoblasts (CTBp), cytotrophoblasts (CTB), pre-syncytiotrophoblasts (STBpre), syncytiotrophoblasts (STB), DUX4-high cells, or HSV-1-high cells. Multiple clusters from a single cell type are annotate with a -number after the cluster name. **(B-C)** Dot plots showing the average expression (ALRA assay) and proportion of cells expressing selected DSGs (B) and HSV-1 transcripts (C) in each cluster separated into mock-infected samples (top) and HSV-1-infected samples (bottom). **(D)** Pseudobulk analysis identifying significant DEGs (Log2(fold-change) > 1 and adjusted *p*-value < 0.05) between HSV-1-infected samples and mock-infected samples. Results from all clusters other than the DUX4-high and HSV-1-high clusters are shown. Genes that belong to the DUX4.101 list are displayed as red circles and HSV-1 transcripts are displayed as blue circles. Also see **Table S6**. **(E)** UMAP of re-clustered cells from HSV-1 infected samples with >0.1% HSV-1 reads. Clusters are labeled as CTBp, CTB, preSTB/STB, DUX4-high, or HSV-1-high. Multiple clusters from a single cell type are annotate with a -number after the cluster name. Slingshot trajectories are superimposed and numbered. **(F)** Featureplots of the pseudotime values (0-100) assigned to each cell along each trajectory (top and lower left). Featureplot showing the cells that were part of the final 15% of cells in a trajectory and designated as part of a trajectory terminus (lower right). **(G)** Pseudobulk analysis identifying significant DEGs (Log2(fold-change) > 1 and adjusted *p*-value < 0.05) between terminus 1 and terminus 2. Genes that belong to the DUX4.101 list are displayed as red circles and HSV-1 transcripts are displayed as blue circles. **(H-I)** Plots of the average expression (scaled to between 0 and 1) of all HSV-1 transcripts (H) and all DSGs (I) along pseudotime 1 and pseudotime 2. Relative peak expression of each HSV-1 transcript (H) and each DSG (I) along each trajectory is shown in a plot to the right. **(J-K)** Featureplots showing percent HSV-1 reads (J) and the expression of the DSG *TRIM49* (K).

We first investigated which cells expressed DUX4/DSG and HSV-1 transcripts. We found that *DUX4* and DSGs were only expressed in HSV-1-infected samples, with one CTB cluster exhibiting particularly high levels of expression (termed “DUX4-high”). *DUX4* and DSG expression was also high in the STB cluster, consistent with the results observed in the trophoblast stem cell model **(Figure 6B and Figure S5I)**. In contrast, most HSV-1 transcripts, including all transcripts classified as late genes (γ1 and γ2) were expressed highest in a distinct CTB cluster (termed “HSV-1-high”) **(Figure 6C and Figure S5J-K)**. A handful of immediate early (α), early (β), and unclassified HSV-1 genes were expressed at comparable or higher levels in the DUX4-high cluster when compared to the HSV-1-high cluster. As HSV-1 genes expression could be observed outside of the HSV-1- and DUX4-high clusters **(Figure 6B-C)**, we next investigated how HSV-1 infection impacted gene expression in these clusters. We performed pseudobulk analysis to define differential gene expression between the HSV-1-infected and mock-infected samples in each of these clusters **(Figure 6D and Table S6)**. Other than HSV-1 genes, DSGs were found to be amongst the most highly differentially expressed genes in all clusters. These data demonstrate that all trophoblast lineages in TOs can respond to HSV-1 infection with the expression of DUX4 and DSGs.

The observation that peak DUX4/DSG expression and peak HSV-1 transcript expression were found in separate HSV-1-infected clusters suggests two possibilities: 1) that DUX4 and DSG expression precedes maximal HSV-1 gene expression during the normal course of replication, or 2) that DUX4 and DSG expression defines a distinct fate of HSV-1-infected cells in which replication is impaired. To determine which was the case, we performed trajectory analysis which seeks to computationally infer the ordered progression of cells along a “pseudotime.” Cells from HSV-1-infected samples with > 0.1% of reads mapping to the HSV-1 genome were re-analyzed independently **(Figure 6E)**. Clustering yielded eight clusters that were defined as described for previous analyzes **(Figure S5G-J)**. Trajectories were predicted using the Slingshot algorithm starting with the CTB-2 cluster and with undefined end clusters.^48^ Three trajectories were inferred, one that leads toward peak DSG expression (pseudotime 1), one that leads toward the peak of HSV-1 gene expression (pseudotime 2), and one that follows differentiation into the STB (pseudotime 3) **(Figure 6F)**. We performed pseudobulk analysis comparing cells at the termini of pseudotime 1 and 2 to determine differential gene expression. We found that all but one HSV-1 transcript was more highly expressed in terminus 2 than 1 **(Figure 6G)**. Conversely, 64 DSGs from the DUX4.101 list were more highly expressed in terminus 1 than 2. Four DSGs followed the opposite pattern, and the remainder were not statistically significant. To visualize gene expression patterns along these trajectories, fitGAM was used to calculate the average expression of HSV-1 transcripts and DSGs along pseudotime trajectories 1 and 2. While HSV-1 transcripts increased along pseudotime 1, most reached only a small fraction of peak expression **(Figure 6H)**. All but two HSV-1 transcripts instead reached maximum gene expression along pseudotime 2. DSGs followed the opposite pattern. Nearly all DSGs reached peak expression along pseudotime 1, while they reached only a fraction of peak expression along pseudotime 2 **(Figure 6I)**. Interestingly, peak DSG expression comes after the rise and fall of HSV-1 genes expression along pseudotime 1. This pattern is borne out by the expression of representative features such as the percent of transcripts mapping to HSV-1, GFP expression from the HSV-1 genome, and expression of DSGs *TRIM49* and *PRAMEF1* **(Figure 6J-K and Figure S5J)**. Taken together, these data suggest that high DSG expression defines a low viral gene expression trajectory in HSV-1-infected trophoblasts.

### DUX4-stimulated genes exert anti-herpesvirus activity

Given that DUX4 and DSGs were highly upregulated in response to herpesviruses and defined a trajectory associated with low viral gene expression following infection, we next sought to determine whether these genes exhibited anti-herpesvirus activity. As the cause of FSHD, many studies have attempted to engineer cells to block or impair DUX4 expression.^49^ In genetically normal cells, hundreds of copies of DUX4 coding sequence are repeated within D4Z4 arrays that are located in the proximal sub-telomeric domains of chromosomes 4 and 10. D4Z4 has additional homologous regions in many other chromosomes.^50^ For this reason, traditional CRIPSR-Cas9 editing within the DUX4 ORF is not generally employed as it would generate many double-stranded breaks, leading to genome instability and off-target effects.^49^ Instead, studies have employed CRISPR interference (CRISPRi),^51^ CRISPR/Cas9-directed base editing,^52^ and dominant-negative DUX4 constructs,^53^ all with varying degrees of knockdown efficiency. We attempted several similar methods of impairing DUX4 activity in TOs including *DUX4*-targeting shRNAs **(Figure S6A-D)**, CRISPRi targeting the transcription start site of the *DUX4* locus **(Figure S6E-G)**, and a dominant-negative DUX4 construct where the transactivation domain was swapped for the KRAB domain of ZIM3 **(Figure S6H-L)**. Although we achieved modest suppression of *DUX4* using shRNAs and of the DSGs *TRIM43* and *PRAMEF1* using the DUX4-ZIM3 construct, DSGs in these contexts were still expressed at ∼250- and 500-fold more than in mock samples (which were at or near the limit of detection). Accordingly, these interventions had insignificant impact on HSV-1 gene expression **(Figure S6D and L)**. For this reason, we decided to instead interrogate the top DSGs induced by herpesviruses in TOs for anti-herpesvirus activity. The biological functions of DSGs are largely unknown. Therefore, we systematically screened individual DSGs for antiviral activity against HCMV and HSV-1 using an overexpression system in cells permissive to virus replication. We selected DSGs that were highly induced in response to iDUX4 expression, and HCMV TB40/E, HCMV AD169r, or HSV-1 infection **(Figure S7A)**. Many DSGs were within transposable elements, causing them to have many orthologues in the human genome which are often other DSGs.^34^ Among DSGs with >90% peptide identity, a single ORF was selected **(Figure S7B)**. The remaining genes were prioritized to a list of 30 based on median log_2_ (fold-change) in TO datasets **(Figure S7C)**. Codon-optimized ORFs of these genes were cloned into lentiviral expression vectors that co-expressed puromycin resistance and Tag2BFP on a polycistronic transcript **(Figure 7A)**. MRC5 cells, which are permissive to HCMV and HSV-1, were arrayed in 96-well plates and transduced with these lentiviruses as well as a control lentivirus (ORF_Stuffer) and a vector expressing a codon-optimized *DUX4* ORF. Cells were selected with 2 μg/mL puromycin for 48 h. Cells were then infected with 5 x 10^4^ pfu / well of HCMV TB40/E (mCherry) or 5 x 10^3^ pfu / well of HSV-1 (GFP-UL25). These infection conditions were chosen to observe a single round of HCMV replication in a high percentage of cells over 96 h, while allowing for multiple rounds of HSV-1 replication over a similar timespan. Replication progression was monitored over time using high-content automated fluorescence microscopy **(Figure 7A)**. Overexpression of six DSGs as well as DUX4 exhibited cell toxicity in MRC5 cells **(Figure S7D-E)**.

**Figure 7.**
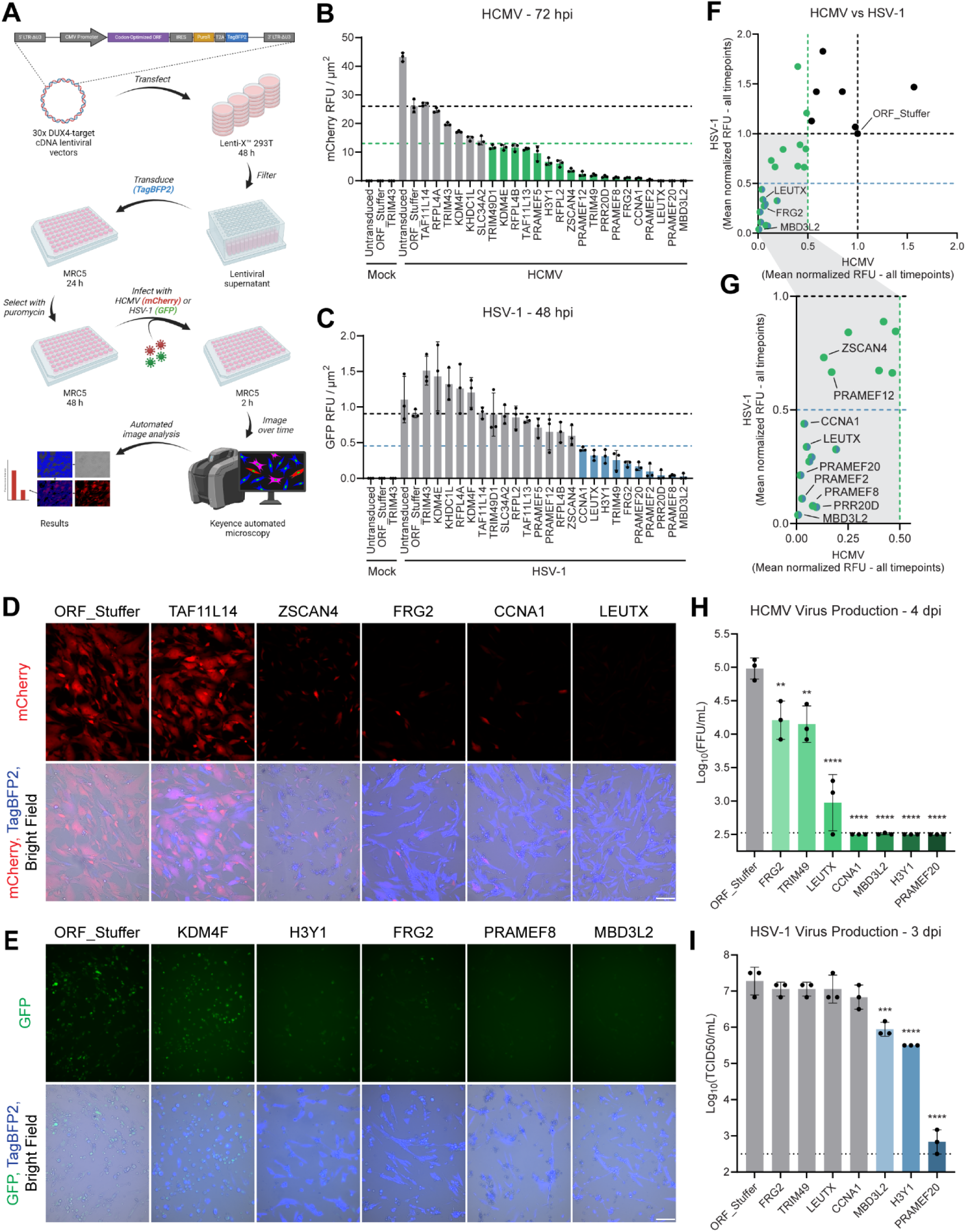
DUX4-stimulated genes exert anti-herpesvirus activity, **(A)** Schematic representing the screening method in MRC5 cells. **(B-C)** Bar graphs depicting representative timepoints for HCMV (B; 72 hpi) and HSV-1 (C; 48 hpi). Black circles represent each individual datapoint. Bars display mean ± SD. **(D)** Representative DSGs for HCMV: one with minimal impact and four top hits. Widefield micrographs of representative fields from 72 hpi. mCherry is portrayed in red in the top row, and merged mCherry (red), TagBFP2 (blue), and brightfield (grayscale) is portrayed in the bottom row. Scale bars = 200 μm. **(E)** Representative DSGs for HSV-1: one with minimal impact and four top hits. Widefield micrographs of representative fields from 48 hpi. GFP is portrayed in green in the top row, and merged GFP (green), TagBFP2 (blue), and brightfield (grayscale) is portrayed in the bottom row. Scale bars = 200 μm. **(F)** Mean normalized relative fluorescence units (RFU) from HCMV in cells expressing each DSG plotted against the mean normalized RFU from HSV-1 in cells expressing each DSG (mean of all timepoints). **(G)** Zoomed in view of panel (F). Black dashed lines in graphs B, C, F, and G represent the mean expression of ORF_Stuffer. Colored dashed lines represent 0.5x the black line. (H-I) Bar graphs depicting virus production from HCMV (H) and HSV-1 (I). HCMV and HSV-1 infectious particles were quantified as focus-forming units (FFU) and 50% tissue culture infectious dose (TCID50) respectively. Black circles represent each individual datapoint. Bars display mean ± SD.Black dotted lines in these graphs represent the limit of detection. Significance was determined by ANOVA with the Tukey multiple comparisons tests. ** p < 0.01, *** p < 0.001, **** p < 0.0001.

Fluorescence intensity of the exogenously expressed tags was quantified, and hits at each timepoint were defined as conditions that reduced the fluorescence intensity to less than half of that of ORF_Stuffer **(Figure S7F)**. We observed 17, 21, 18, and 15 hits for HCMV at 1, 2, 3, and 4 dpi respectively **(Figure 7B and D and Figure S8A-C)**. 13, 10, and 8 hits were observed for HSV-1 at 1, 2, and 3 dpi **(Figure 7C and E and Figure S8E-F)**. Ten genes consistently restricted replication of both viruses: *CCNA1*, *FRG2*, *H3Y1*, *LEUTX*, *MBD3L2*, *PRAMEF2*, *PRAMEF8*, *PRAMEF20*, *PRR20D*, and *TRIM49* **(Figure 7F-G)**. A handful of candidates were found to be antiviral to HCMV but not HSV-1. These included *PRAMFE12* and *ZSCAN4*, both of which consistently reduced mCherry expression from HCMV to less than 25% of ORF_Stuffer while having minimal impact on GFP expression from HSV-1 **(Figure 7G)**. The top hits from both viruses suppressed fluorescence protein expression throughout all timepoints **(Figure S8D and G)**.

Reduction in fluorescent tag expression does not necessarily indicate a measurable decrease in virus output. To explore the magnitude of the impact of DSGs on herpesvirus virion production, MRC5 cells were transduced with seven of the top DSGs or the ORF_Stuffer control vector and selected with puromycin as described above. These cells were then infected with HCMV TB40/E mCherry (MOI of ∼2) or HSV-1 GFP-UL25 (MOI ∼0.1) followed by washing three times with PBS to remove residual inoculum. Media was collected at 4 dpi for HCMV and 3 dpi for HSV-1 allowing for a single round of replication or multiple rounds of replication respectively. Quantification of virus in the supernatant revealed that all seven DSGs significantly reduced HCMV virus production with many of DSGs yielding virus concentrations near the limit of detection **(Figure 7H)**. Consistent with the results from the screen, the effect of DSGs on HSV-1 replication was less pronounced compared to HCMV **(Figure 7I)**. However, MBD3L2, H3Y1, and PRAMEF20 reduced HSV-1 virus production by between one and four logarithms. These data show that many DSGs function as antiviral mediators.

## Discussion

Placental trophoblasts form the barrier between the maternal and fetal compartments of the MFI and possess cell-autonomous mechanisms of protecting the developing fetus from the vertical transmission of viruses.^1,2^ Furthermore, trophoblasts have been shown to restrict the replication of prominent teratogenic herpesviruses such as HCMV.^20,21^ In this study, we profiled the response of TOs to seven teratogenic viruses and identified the expression of DUX4 and downstream target DSGs as a unique response to herpesvirus infections. Comparison between TOs and matched maternally-derived DOs revealed that DUX4 and DSGs are induced to significantly higher levels in TOs by both HSV-1 and HCMV, suggesting that it is uniquely induced in trophoblasts. Using scRNAseq and pseudotime trajectory analysis, we identified a unique pathway in HSV-1-infected TOs. This trajectory was distinct from the typical virus replication trajectory and was characterized by the expression of *DUX4* and downstream target DSGs, consistent with this program being antiviral. Overexpression screening of the most highly induced DSGs demonstrated that many of these genes significantly impaired the replication of HSV-1 and HCMV in permissive cells. These findings uncover a previously unrecognized antiviral pathway in human trophoblasts, wherein the induction of DUX4 and its downstream target genes serves as a cell-intrinsic defense mechanism against herpesvirus infections. This discovery highlights the unique antiviral strategies employed by trophoblasts to protect the developing fetus from teratogenic viral threats.

A relationship between the DUX4 program and herpesvirus infection has emerged, but there is not a consensus on its role in virus replication. Expression of DUX4 or select DSGs have been observed in response to infection with alpha, beta, and gamma herpesviruses in cell culture as well as in herpesvirus-mediated cancers.^37,54–56^ Induction of DUX4 following herpesvirus infection was originally described based on the antiviral activity of the E3 ubiquitin ligase TRIM43.^37^ TRIM43 was proposed to impair herpesvirus replication by directing components of the centrosome for degradation leading to destabilization of the nuclear envelope.^37^ Recently, knockdown of the DSGs *RASD1* and *RRAD*, which are Ras-related GTPases, was found to augment HSV-1 virion production in fibroblasts.^38^ As the above DSGs exert antiviral activity, DUX4 and DSGs have been suggested to be a part of the cell-intrinsic defenses against herpesvirus infection. Contrary to this model, a recent study has shown that HSV-1 and HSV-2 replication is impaired in DUX4 KO HAP1 cells and DUX4 knockdown HEK293 cells, suggesting that aspects of the DUX4 program may be coopted by the virus to facilitate replication in some contexts.^56^ In a primary cell-derived trophoblast system, we found that the expression of DUX4 and the majority of DSGs defined a low viral gene expression trajectory following HSV-1 infection in TOs. Consistent with this, overexpression of the majority of DSGs restricted HSV-1 and/or HCMV replication in MRC5 fibroblasts. Thus, our data support a model in which DSGs are a component of the cell-intrinsic defenses against herpesvirus replication. In the cell types examined in previous studies, DSG induction typically occurs alongside canonical IFN antiviral signaling, which likely serves as the primary antiviral defense mechanism. In contrast, our findings reveal that in human trophoblasts, the DUX4 pathway operates as the predominant defense mechanism against herpesvirus infections, representing a specialized antiviral strategy unique to this cell type. Our data expands our conception of DUX4 and DSGs from isolated antiviral genes controlled by a common transcription factor to a coordinated cell-intrinsic antiviral pathway in human trophoblasts.

Most DSGs are not expressed at detectable levels in normal adult cells, and many remain largely uncharacterized. Their specific roles and potential functions are poorly understood, highlighting a significant gap in our knowledge about these genes and their biological importance. Apart from *TRIM43*, *RASD1*, and *RRAD*,^37,38^ it remained unclear how many DSGs exert anti-viral activity. Thus, we tested the impact of overexpression of 30 DSGs that were highly expressed in TOs on HSV-1 and HCMV replication in permissive MRC5 cells. Twenty-one of these genes reduced fluorescent protein expression of at least one virus by >50% during at least one timepoint. If we extrapolate to include genes that were excluded based on the >90% peptide identity cutoff, this screen identified 42 novel potentially antiviral human ORFs. Of note, one of the three genes that had little impact on the replication of either virus at any timepoint was *TRIM43*. *RASD1* and *RRAD* were not tested in this screen because they did not meet the expression cutoffs in TOs for inclusion. Additionally, none of the DSGs in the screen led to an increase in viral fluorescent protein expression above that of the control vector. One of the top antiviral DSGs identified by the screen was *CCNA1*, which is a meiosis-specific cyclin A gene in mammals.^57^ Although, *CCNA1* has not been previously studied in the context of virus replication, its paralog *CCNA2* restricts HCMV replication.^58^ The largest group of genes represented in our screen were the PRAME (<u>pr</u>eferentially expressed <u>a</u>ntigen in <u>me</u>lanoma) family of leucine-rich repeat proteins. All five of the PRAME family genes tested displayed some level of anti-viral activity, with *PRAMEF2*, *PRAEMF8*, and *PRAMEF20* being among the top hits for both viruses. The only member of this family in humans to have been studied mechanistically is *PRAME*, which can function as a negative regulator of retinoic acid signaling.^59,60^ However, *PRAME* does not appear to be a DSG based on our analyses, and involvement in retinoic acid signaling has not been confirmed with any other PRAME family members which share at most 53% peptide identity with *PRAME*. The PRAME family is extensively duplicated in mammals. Additionally, both the PRAME family and *LEUTX*, another top hit from this screen, have high allelic variation in humans and exhibit signs of positive selection, which are common features of antiviral gene families.^61–63^ Our findings suggest that herpesvirus infections may have exerted evolutionary selective pressure on the development and retention of these genes. We conclude that many of the highly upregulated DSGs in trophoblasts possess strong anti-herpesvirus activity, positioning them as compelling targets for future mechanistic studies to better understand their roles in antiviral defense and potential therapeutic applications.

A detailed mechanism of how herpesvirus infection triggers *DUX4* expression has yet to be reported. A recent study suggests that the HSV-1 immediate early genes RL2 (ICP0) and US1 (ICP4) are necessary and sufficient to induce DUX4 expression in HSV-1 infection.^56^ In our scRNAseq data of HSV-1 infection in TOs, we observed that 9 out of 50 HSV-1 genes were expressed at comparable or higher levels in the DUX4-high cluster when compared to the HSV-1-high cluster. All of these genes were immediate early (α), early (β), or unclassified HSV-1 transcripts. Thus, DUX4 expression being triggered by the activities of one or several immediate early genes may explain this expression pattern. Importantly, US1 was not among the HSV-1 genes with higher expression in the DUX4-high cluster and the expression of RL2 was too low to accurately conclude a pattern of expression between these clusters. Another observation that may point to how DUX4 is controlled is that TOs respond to HCMV and HSV-1 with much higher expression of both *DUX4* and downstream DSGs than matched maternal DOs. As trophoblasts uniquely constitutively secrete IFN-λs,^12,18–20^ we hypothesized that constitutive type-III IFN signaling may modulate *DUX4* and DSG expression. However, knock-out of *IFNLR1* did not decrease *DUX4* or DSG expression in TOs following HSV-1 infection, demonstrating that constitutive type III IFN signaling does not influence this pathway. Nonetheless our data indicate that TOs respond to herpesvirus infection with uniquely high expression of *DUX4* and DSGs, which could be used in future studies to investigate how this pathway is induced and modulated during herpesvirus infection.

Immune responses at the MFI must be tightly regulated to prevent damage to the developing fetus. For example, in mice the type I IFN response, either induced by viruses or poly I:C, can cause placental damage and lead to fetal demise.^14–16,64^ Although RuV infection induced IFN and ISG expression in TOs, demonstrating that IFN pathways can be activated in this model, HSV-1, HSV-2, and HCMV infection had minimal impact on the expression of IFNs and ISGs. The induction of DSGs may also have potential negative impacts on trophoblast function, raising the possibility of trade-offs between antiviral defense and maintaining normal cellular processes. Expression of DUX4 and six of the DSGs that were tested in our screen led to cell death in MRC5 cells, suggesting that expression of these genes could also be lethal in trophoblasts. Additionally, although scRNAseq analysis of iDUX4 expression and HSV-1 infection revealed that all trophoblast cell types can produce DSGs, the STB was one of the clusters that expressed the highest levels of DSGs in response to HSV-1 infection. The STB is a critical trophoblast cell type that participates in placental hormone secretion and nutrient, gas, and waste exchange. The impacts of expression of a potently lethal gene in a polynuclear fused cell such as the STB are unclear. DUX4 causes FSHD when aberrantly expressed in adult muscle cells—another polynuclear fused cell type.^35,36^ Finally, many DSGs are small molecule transporters, such as the sodium phosphate transporter *SLC34A2* and the amino acid transporter *SLC36A2,*^65,66^ which were induced by 240-fold and 11-fold respectively by iDUX4 expression in TOs. In MRC5 cells expression of SLC34A2 had a small impact on HCMV replication at one timepoint and no impact on HSV-1 replication. However, an increase in expression of small molecule transporters could disrupt the crucial functions of nutrient and waste exchange between the maternal and fetal blood supplies across the STB. Thus, our data present the possibility that DUX4 expression in trophoblasts following herpesvirus infection could contribute to impaired placental function.

The data presented here establish that DUX4 and its transcriptional targets, termed DSGs, form the central response mechanism of trophoblasts to herpesvirus infection, underscoring their critical role in the antiviral defense of these specialized cells. Like ISGs, DSGs are induced as part of an antiviral defense mechanism and encode non-redundant effectors that likely target distinct aspects of viral replication. However, the DUX4-DSG pathway appears uniquely adapted for pregnancy, enabling a cell-intrinsic antiviral response in trophoblasts without activating paracrine signaling that could disrupt maternal immune tolerance. Given the high prevalence of herpesvirus exposure during pregnancy, this pathway may have evolved to balance effective antiviral defense with the preservation of placental function and immune tolerance, ensuring optimal pregnancy outcomes. These insights not only deepen our understanding of trophoblast biology but also open new avenues for exploring therapeutic strategies to protect against teratogenic viral infections.

## Acknowledgements

This work was supported by NIH grants F32-AI183633 (JH), R01-AI173333 (CC), P01-AI129859 (CC), and R01-AI145828 (CC). We would like to acknowledge Phillip Kolb (University Hospital Freiburg), Rebecca Göttler (University Hospital Freiburg), Nat Moorman (University of North Carolina at Chapel Hill) and Bekah Dickmander (University of North Carolina at Chapel Hill) for assistance with HCMV propagation.

## Materials and Methods

### Culture of trophoblast and decidua organoids

TO and DO lines employed in this study were derived from full-term placental tissue as previously described.^20,67,68^ Human tissue used in this study was obtained from Duke University after approval from the Duke University Institutional Review Board and in accordance with the guidelines of Duke University human tissue procurement. TOs were cultured by plating in domes of Matrigel (Corning 356231) and maintained with term trophoblast organoid medium (tTOM) [Advanced DMEM/F12 (Life Technologies, 12634010) supplemented with 1X B27 (Life Technologies, 17504044), 1× N2 (Life Technologies, 17502-048), 10% FBS (vol/vol, Cytiva HyClone, SH30070.03), 2 mM GlutaMAX^TM^ supplement (Life Technologies, 35050061), 100 µg/ml Primocin (InvivoGen, ant-pm-1), 1.25 mM N-acetyl-L-cysteine (Sigma, A9165), 500 nM A83-01 (Tocris, 2939), 1.5 µM CHIR99021 (Tocris, 4423), 50 ng/ml recombinant human EGF (Gibco, PHG0314), 80 ng/ml recombinant human R-spondin 1 (R&D systems, 4645-RS-100), 100 ng/ml recombinant human FGF2 (Peprotech, 100-18C), 50 ng/ml recombinant human HGF (Peprotech, 100-39), 10 mM nicotinamide (Sigma, N0636-100G), 5 µM Y-27632 (Sigma, Y0503-1MG) and 2.5 µM prostaglandin E2 (PGE2, Tocris, 2296)].^20^ DOs were maintained in Expansion Medium (ExM) [Advanced DMEM/F12 (Life Technologies, 12634010) supplemented with 1X B27 without vitamin A (Life Technologies, 12587010), 1× N2 (Life Technologies, 17502-048), 2 mM GlutaMAX^TM^ supplement (Life Technologies, 35050061), 100 µg/ml Primocin (InvivoGen, ant-pm-1), 1.25 mM N-acetyl-L-cysteine (Sigma, A9165), 500 nM A83-01 (Tocris, 2939), 50 ng/ml recombinant human EGF (Gibco, PHG0314), 100 ng/mL recombinant human Noggin (Peprotech, 120-10C), 80 ng/ml recombinant human R-spondin 1 (R&D systems, 4645-RS-100), 100 ng/ml recombinant human FGF-10 (Peprotech, 100-26), 50 ng/ml recombinant human HGF (Peprotech, 100-39), 10 mM nicotinamide (Sigma, N0636-100G), and 2.5 µM Y-27632 (Sigma, Y0503-1MG)].^68^ Organoid cultures were maintained in a humidified incubator with 5% CO_2_ at 37°C, and medium was renewed every 2-3 days. After 4-5 days in culture, organoids were grown in their respective growth media without Y-27632. Organoids were passaged after 5-7 days in culture by collecting Matrigel domes, digesting with TrypLE Express (Gibco, 12605-028) for 6-12 min at 37°C, then mechanically dissociating into single cells and small fragments by manual pipetting. Resulting cells were resuspended in Matrigel and plated as domes in 24-well tissue culture plates.

### Culture of cell lines and viruses

Lenti-X™ 293T cells (Takara, 632180), ARPE-19 cells (ATCC, CRL-2302) and Vero (ATCC, CCL-81) cells were cultured in Dulbecco’s minimal essential medium (DMEM) with 10% FBS and 1% penicillin/streptomycin (Gibco, 15140). MRC5 cells (ATCC, CCL-171) were cultured in DMEM with 10% FBS, 1% penicillin/streptomycin (Gibco, 15140), 1x HyClone™ non-essential amino acids (Cytiva, SH30238.01), 1 mM sodium pyruvate (Cytiva, SH30239.01). Cell lines were tested for mycoplasma using the MycoStrip Mycoplasma Detection Kit (Invivogen, rep-mys-50).

mCherry-tagged HCMV TB40/E (a gift from N. Moorman^69^) was propagated by infecting ARPE-19 (MOI ≅ 0.01) followed by extended incubation until >90% of cells showed CPE. Growth on ARPE-19 cells was chosen to select for epithelial tropism. Both supernatant and cells were harvested, cells were gently lysed by freeze/thaw, and pooled cell-associated and cell-free virus was concentrated on a 20% sucrose cushion (20,000 rpm). Virus pellet was the resuspended in sterile media before titration on MRC5 cells and ARPE-19 cells. The remaining virus stocks were previously prepared, and aliquots were pulled from storage at -80°C for experiments. GFP-UL25 HSV-1 KOS strain (a gift from F. Homa^70^),^11^ HSV-2 333 strain (a gift from H. Lazear), YFP-tagged VACV Western Reserve strain (a gift from S. Cherry),^11^ RuV M33 strain (ATCC, VR-315),^71^ and ZIKV Paraiba-2015 strain (a gift from D. Watkins)^12^ were all titrated on Vero cells. Echo5 Noyce strain was titrated on HeLa cells ^72^.

### Virus infection in organoids

Virus infections in organoids were performed either on coat-seeded cultures or in suspension. Matrigel coats were generated by covering the bottom of a 48-well plate or 8-well chamber slide with 20 uL of undiluted Matrigel, which was then allowed to polymerize for ∼10 min. Dissociated organoids were seeded on Matrigel coats following passage and allowed to grow in for 5-7 days prior to infection. The coat-culturing method allows organoids to grow into 3D spherical structures, but leaves the majority of the surface of each organoid unit exposed to the media rather than fully imbedded in matrigel as is typical in Matrigel domes. Coat-seeded organoids were infected by replacing the culturing media with inoculum (virus diluted in tTOM), incubating for 2 h at 37°C, and then replacing the inoculum with fresh culturing media. Suspension infections were performed by first releasing organoids from Matrigel domes using cell recovery solution (Corning, 354253) at 4°C for 20-30 min. Organoids were pelleted by gravity, washed with PBS, and pelleted again. Inoculum was then added to the released organoids. Infections were allowed to proceed for 2 h at 37°C, agitating the culture every 30 min to keep the organoids from aggregating. Organoids were then pelleted, resuspended in Matrigel, seeded in domes in 24-well plates, and submerged with normal tTOM. Prior to infections, parallel wells of organoids were disrupted and counted to calculate MOI. MOI for each experiment is specified in the results and figure legends.

### Lentivirus production

Lentiviral particles were produced in Lenti-X™ 293T cells (Takara, 632180) by transfection of a lentiviral transfer plasmid along with psPAX2 (Addgene #12260) and pMD2.G (Addgene #12259) packaging plasmids using TransIT®-293 transfection reagent (Mirus Bio, MIR 2700). 293T cells were refed with intended target cell medium at 24 h post transfection. Supernatant was collected at 48 h post transfection, filtered with 0.45 μm filter, then stored at -80°C prior to transduction of target cells. psPAX2 and pMD2.G were gifts from Didier Trono. pCW57.1-DUX4-CA, used to generate iDUX4 lentiviral particles, was a gift from Stephen Tapscott (Addgene #99281).^45^ Lentiviral plasmids for the overexpression of DSGs, CRISPR-Cas9-mediated editing of IFNLR1, shRNAs, CRISPRi, and dominant-negative DUX4 overexpression were constructed by VectorBuilder. Descriptions of these vectors along with vector IDs (that can be used to retrieve detailed information on them) can be found in **Table S7**.

### Transduction and single-cell cloning of trophoblast organoids

Trophoblast organoids were transduced by first dissociating them into single-cell suspension following the same protocol for passaging described above. The dissociated single cell suspension was then incubated with filtered lentiviral supernatant and 8 μg/mL polybrene overnight in an ultralow attachment plate at 37°C overnight. The transduced cell suspension was resuspended in Matrigel and plated as domes. After 2-3 days of recovery, transduced cells were selected for by supplementing the media with 2 ug/mL puromycin for 4-5 days. Single cell clones of CRISPR-Cas9-edited organoids were generated by isolating and propagating individual organoid units one passage after selection. This process takes advantage of the likelihood that most organoids in Matrigel arise from a single or very few starting stem cells. Clonal organoid lines were screened for knockout by isolating genomic DNA using GenElute Mammalian Genomic DNA Purification Kit (Sigma, G1N350), amplifying the sgRNA target region using high-fidelity PCR, and the sequencing the purified PCR product with sanger sequencing at GENEWIZ. The resulting capillary electrophoresis traces were analyzed using the ICE analysis tool from Synthego. The primers used to amplify the sgRNA target editing site in *IFNLR1* are listed in **Table S7**.

### Immunofluorescence, confocal microscopy, and confocal image analysis

For immunofluorescence staining, organoids cultured in a layer of matrigel on 8-well chamber slides. Organoids were fixed with 4% paraformaldehyde for 30 min, permeabilized with PBS with 0.5% Triton X-100 for 60 min, then blocked with 10% normal goat serum in PBS with 0.05% TWEEN-20 for 60 min. Organoids were then incubated with primary antibody diluted in blocking solution at 4°C overnight. Organoids were then washed with PBS and then incubated for 2 hr at room temperature with Alexa Fluor-conjugated secondary antibodies. Organoids were washed again with PBS and mounted in Vectashield (Vector Laboratories) containing 4′,6-diamidino-2-phenylindole (DAPI). The following antibodies were used for staining: DUX4 (Sigma, MABD116), anti-V5-tag (Cell Signaling Technologies, #58009), Alexa Fluor 488 Goat anti-Mouse IgG secondary antibody (Invitrogen, A11029), and Alexa Fluor 594 Goat anti-Rabbit IgG secondary antibody (Invitrogen, A11037). Images were captured on an Olympus Fluoview FV3000 laser scanning confocal microscope.

Nuclear mean gray value (MGV) measurements were calculated using an ImageJ macro that first runs the StarDist package on images from the DAPI channel to generate individual regions on interest (ROIs) for every nucleus.^73^ These ROIs are then applied to images from other channels to measure there MGVs within each nucleus.

### Overexpression screen and automated microscopy

Lentivirus particles for constructs used in the screen were prepared as described above and arrayed into 96-well plate format for this screen. MRC5 cells were seeded into 96-well plate format at a density of 5 x 10^3^ cells per well. The following day, the media on these cells was replaced with MRC5 culturing medium supplemented with 16 μg/mL polybrene. 100 μL of the appropriate lentivirus or mock media was added to each well. Transductions were performed in triplicate. Transduction plates were then incubated statically at 37°C for 30 min, centrifuged at 800xg for 30 min, then incubated for an additional hour statically at 37°C. Transduction media was then replaced with fresh culturing media. 24 h post transduction, the media was replaced with MRC5 culturing media supplemented with 2 μg/mL puromycin, and selection was allowed to proceed for 48 h. 72 h post transduction, selected cells were infected with HCMV TB40/E mCherry (5 x 10^4^ particles/well), HSV-1 KOS GFP (5 x 10^3^ particles/well), or mock media. Inoculums were applied to cells for 2 h statically at 37°C. Inoculum was then replaced with fresh media. A Keyence BZ-X810 all-in-one automated microscope was used to capture fluorescent and bright field images at 10x magnification of every well at 24 h intervals for the duration of each experiment. Five fields of images were captured per well. Cultures were maintained in a humidified, 37°C chamber during imaging. Lentiviral constructs that led to complete cell death were excluded from further analysis. This was defined by visual morphological assessment of bright field images (spherical cell body and/or detachment) at 0 hpi and 24 hpi, which are timepoints at which CPE is not observed by either virus in control conditions. Fluorescence quantification was performed using BZ-X800 Image Analyzer (Keyence). Cell body areas were defined algorithmically using brightfield images. Fluorescence intensity of the virally encoded tags were integrated within these areas and then normalized to the total cellular area (in μm^2^), yielding an RFU / μm^2^ measurement for each well at each timepoint.

### Virus titration assays

HCMV infectious virus was quantified by focus-forming assay. Briefly, supernatants from HCMV replication experiments were serially diluted, and used to infect MRC5 cells. At 4 dpi, these cells were fixed in 2% paraformaldehyde for 1 h. Cells were then stained for HCMV gene expression by permeabilizing with PBS containing 0.5% Triton X-100 and 1% BSA for 20 min. Cells were then incubated overnight at 4°C with an anti-UL44 primary antibody (Virusys, CA006-100) at a 1:5000 dilution in PBS containing 0.1% TWEEN20 and 1% BSA. Cells were then washed and counter-stained for > 1 h at room temperature with anti-mouse IgG-HRP (Bio-Rad, #1706516) diluted 1:2000 in PBS containing 0.1% TWEEN20 and 1% BSA. Cells were then incubated with DAB (3,3’Diaminobenzidine) chromogen (Abcam, ab64238) following the manufactures instructions. UL44-positive cells at the most dilute sample with detectable staining were counted and used to calculate focus-forming units (FFU) per mL (FFU/mL). HSV-1 infectious virus was quantified using the 50% tissue culture infectious dose (TCID50) method. Briefly, supernatants from HSV-1 replication experiments were serially diluted and used to infect Vero cells. At 7 dpi, cells were fixed and stained simultaneously with a crystal violet solution in PBS containing 2% PFA. The number of wells with detectable monolayer disruption at each dilution were counted and used to calculate TCID50/mL.

### RNA extraction, qRT-PCR, and bulk RNAseq

RNA for qRT-PCR and bulk RNAseq was purified using the Sigma GenElute total mammalian RNA miniprep kit or the Qiagen RNAeasy Mini Kit following the manufacturer’s instruction. cDNA for qRT-PCR was generated from total RNA using iScript cDNA synthesis kit (Bio-Rad) following the manufacturer’s instructions. Quantitative PCR was performed using the iQ SYBR Green Supermix (Bio-Rad, 1708882) on a CFX96 Touch Real-Time PCR Detection System (Bio-Rad). Gene expression was determined based on a ΔCT method normalized to actin. Primers used for qPCR can be found in **Table S7**.

Bulk RNAseq libraries were prepared by the Duke Center for Genomic and Computational Biology using the KAPA mRNA HyperPrep Kit (Roche). Sequencing was performed as follows: on a NovaSeq X Plus using 100 bp paired end sequencing for the diverse virus TO infection experiment, on a NovaSeq 1000 using 100 bp paired end sequencing for the iDUX4 experiment, and on a NextSeq 500 using 75 by paired end sequencing for the HSV-1 infection in matched TO and DO experiment. The following datasets were downloaded from Sequence Read Archive (SRA) and used in analyses: TO/DO infections with HCMV TB40/E and HCMV AD169r (PRJNA869760),^20^ TSC infections with HCMV TB40/E (PRJNA1045985),^21^ iDUX4 expression in iPSCs (PRJNA377314),^34^ and iDUX4 expression in MB135 cells (PRJNA338591).^45^

RNAseq reads were aligned to the human genome (GRCh38) and the appropriate viral genomes for each experiment using QIAGEN CLC Genomics Workbench (v24). Viral genomes used in mapping are as follows: HCMV TB40/E (GenBank: MW439039.1), HSV-1 KOS (GenBank: KT899744.1), HSV-2 333 (GenBank: LS480640.1), VACV WR (GenBank Assembly: GCA_000860085.1), Echo5 Noyce (GenBank: AF083069.1), RuV M33 (GenBank: OM816674.1), and ZIKV Paraíba (GenBank: KX280026.1). Differential gene expression analysis was performed using DESeq2 package in R accounting for multiple donors in the statistical model.^39^ PCA analysis was performed using PCAtools package in R following correction for donor, experimental set, and dpi batch effects using the ComBat function of the sva package.^74^ Heatmaps were generated using log_2_ normalized counts (an output from DESeq2) and the pheatmap package in R. Raw data from the bulk RNAseq experiments reported here have been deposited into SRA (PRJNA1213838).

### Dissociation of organoids for single cell RNAseq

Matrigel domes of organoids were resuspended into media [Advanced DMEM/F12 (Life Technologies, 12634010) supplemented with 1% penicillin/streptomycin (Gibco, 15140), 10 mM HEPES (Gibco, 15630-106), and 2 mM GlutaMAX^TM^ supplement (Life Technologies, 35050061)] and organoids were pelleted by centrifugation. The media was replaced by TrypLE Express (Gibco, 12605-028), and organoids were digested at 37°C for 12 min. Organoids were pelleted again and resuspended in 200 uL resuspension media and vigorously disrupted by manual pipetting (∼180 pipettes per min) for 3 min. The volume of the cell suspension was increased to 1 mL with PBS supplemented with 1% FBS. The cell suspension was then filtered through a 40 μm filter cell strainer (Corning, 352098) using an additional 3 mL of PBS supplemented with 1% FBS to wash the filter. Cells were pelleted by centrifugation and resuspended in 100 μL of PBS supplemented with 1% FBS. Cells were counted and viability tested using trypan blue and automated cell counting (Biorad TC20 automated cell counter). All samples had >1 x 10^6^ live cells/mL and viabilities >70%.

### Single-cell RNAseq library preparation and data analysis

Libraries were prepared from ∼10,000 cells using 10x Genomics Single Cell 3’ reagent kits (v2 chemistry, #CG00052) on a Chromium X instrument (10x Genomics) by the Molecular Genomics Core at the Duke Molecular Physiology Institute. Sequencing was performed on a NovaSeq X sequencer using a S4 flow cell which was predicted to yield around 71k reads / cell. Custom genomes were constructed using the CellRanger package (v6.1.2, 10x Genomics) in Linux by adding either the iDUX4 lentiviral vector sequence or the HSV-1 KOS genome (GenBank: KT899744.1) to the human genome (GRCh38) as additional chromosomes. HSV-1 genes that share a polyadenylation site (e.g. polycistronic or alternatively spliced transcripts) were grouped into single features as they cannot be distinguished by the employed library preparation chemistry. Post-processing and read alignment to these custom genomes were performed using the 10x CellRanger package hosted through the 10x cloud analysis platform. Gene expression matrices were analyzed using the Seurat (version 4) package in R.^75^ Cells with between 2000 and 12000 uniquely expressed genes and between 4% and 25% mitochondrial reads were included for downstream analysis. To eliminate batch effects, samples from different donors within the same condition were normalized and integrated using the SCTransform and FindIntegrationFeatures functions in Seurat while simultaneously regressing the following variables: percent mitochondrial reads, percent ribosomal reads, percent X chromosome reads, percent Y chromosome reads, number of features per cell, number of counts per cell, and percent HSV-1 reads (for the HSV-1 infection dataset) or DUX4/DSG score (for the iDUX4 dataset). The RNA assay was scaled and normalized and ALRA-mediated imputation of the data was performed before subsequent analyses.^76^ Samples from separate conditions were then merged together using the Harmony package in R.^77^ Dimensional reduction was performed using the RunPCA function followed by the RunUMAP function using the top 50 principle component dimensions. The FindNeighbors and FindClusters commands were used to perform clustering at a resolution of 0.6. Determination of an optimal resolution was aided by the clustree package.^78^ Correlation between iDUX4 and all other genes was performed using the Pearson correlation setting of the base cor() function in R. Differential gene expression analysis was performed by retrieving the raw counts from each cluster separated by original sample using the AggregateExpression function and then performing DESeq2 analysis accounting for multiple donors in the statistical model.^39^ HSV-1 transcripts were classified as described in Dremel et al.^79^

To perform trajectory analysis of HSV-1 infected cells, the data were reanalyzed to only include infected cells and to allow these cells to be ordered based on HSV-1 gene expression. Cells with between 2000 and 12000 uniquely expressed genes, >10000 total reads, between 4% and 25% mitochondrial reads, and >0.1% HSV-1 reads were included for downstream analysis. To eliminate batch effects, samples from different donors were normalized and integrated using the SCTransform and FindIntegrationFeatures functions in Seurat while simultaneously regressing percent X chromosome reads, percent Y chromosome reads, and number of counts per cell. Dimensional reduction was performed using the RunPCA function followed by the RunUMAP function using the top 20 principle component dimensions. The FindNeighbors and FindClusters commands were used to perform clustering at a resolution of 0.6. The SlingShot package in R was used to predict pseudotime orders of infected cells from clusters identified in Seurat using a defined starting cluster and undefined end clusters.^48^ The fitGAM function of the tradeSeq package in R was used to fit variable features, HSV-1 genes, and DSGs to a generalized additive model along the predicted pseudotime trajectories.^80^ The predictSmooth function was used to plot the average expression of genes in cells along trajectories. Raw data from the scRNAseq experiments reported here have been deposited into SRA (PRJNA1213838).

### Statistics and reproducibility

All primary organoid-based experiments in this report have been replicated using independent samples from organoid lines derived from multiple donors. Experiments in cell lines and cloned organoids lines were performed with triplicate independent samples. Data in bar charts are presented as mean ± SD. In all graphics, each individual data point is shown when possible. Statistical significance was determined as described in the figure legends. All statistical analyses were performed using GraphPad Prism, unless otherwise specified (e.g. bulk RNAseq and scRNAseq). Unless otherwise specified, p values of <0.05 were considered to be statistically significant.

---

**Figure S1.**
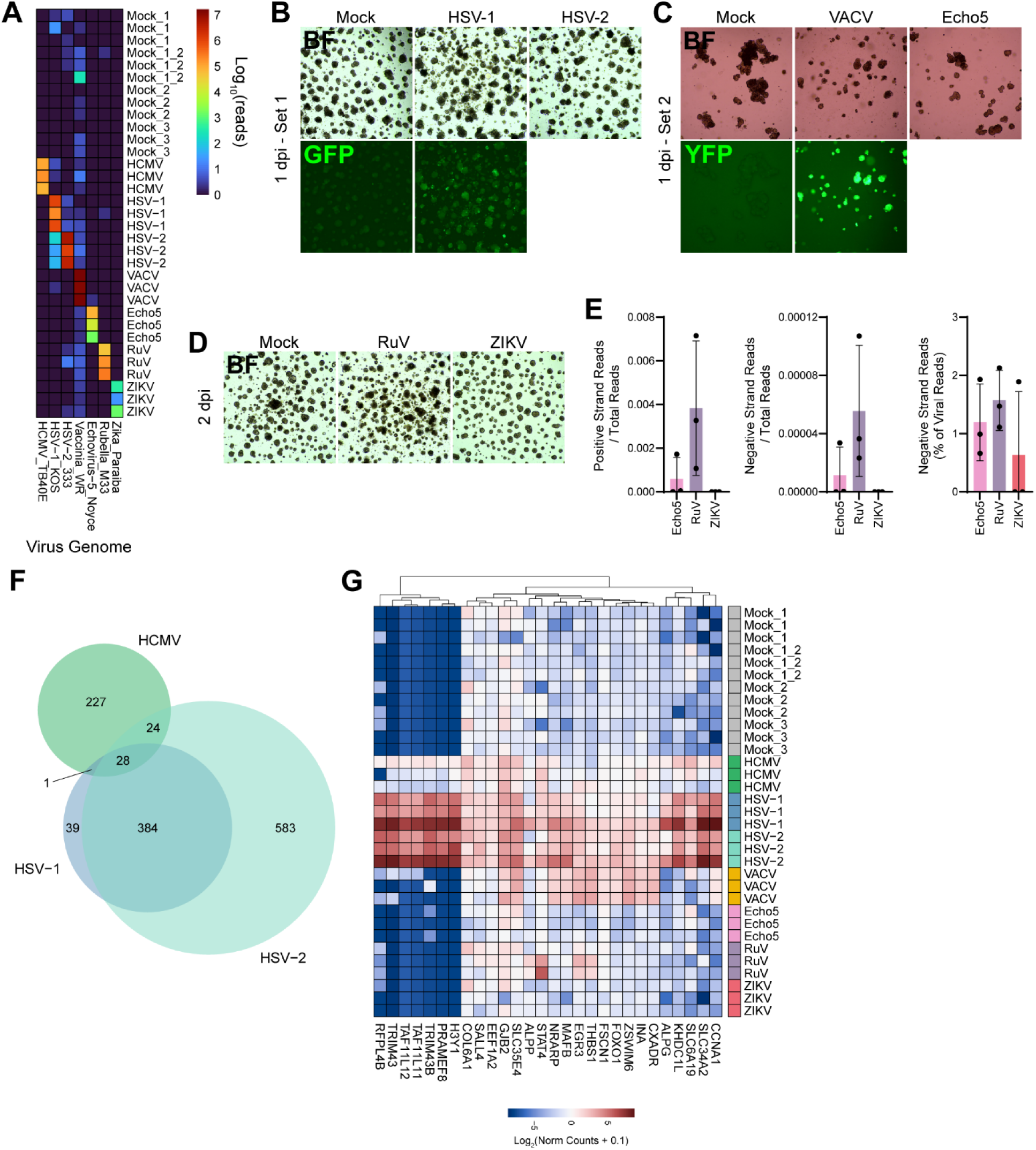
Trophoblast organoids respond distinctly to diverse viruses. Trophoblast organoids from three donors were infected with HCMV, HSV-1, HSV-2, VACV, Echo5, RuV, and ZIKV at an MOI of 2. VACV and Echo5 infections were performed in suspension on organoids released from matrigel that were re-imbedded in matrigel following inoculation. All other infections were performed on spherical organoids grown on matrigel coats. mRNA was purified and analyzed by RNAseq. **(A)** Heatmap portraying the log_10_ reads that map to each of the viral genomes. **(B-D)** Microscopic images of mock and infected organoids. Bright field (BF) images are in the first row of each panel. Fluorescence images were included for HSV-1 (B) and VACV (C). **(D)** Bar plots of the number of reads mapping to the positive and negative strand of the Echo5, RuV, and ZIKV genomes normalized to the total reads in the sample (left two graphs). Bar plot of the percent of viral reads that mapped to the negative strand of the viral genome (right graph). Bars display mean ± SD. Black dots represent each individual data point. **(F)** Venn diagram denoting the number of shared DEGs between each of the herpesvirus infection conditions. **(G)** Heatmap portraying the genes that are induced by all three herpesviruses in TOs. Values are log_2_ normalized counts from DESeq2 analysis plus a pseudo count of 0.1. Key is at the bottom. Red indicates higher expression and blue indicates lower expression. Hierarchical clustering of genes is shown at the top.

**Figure S2.**
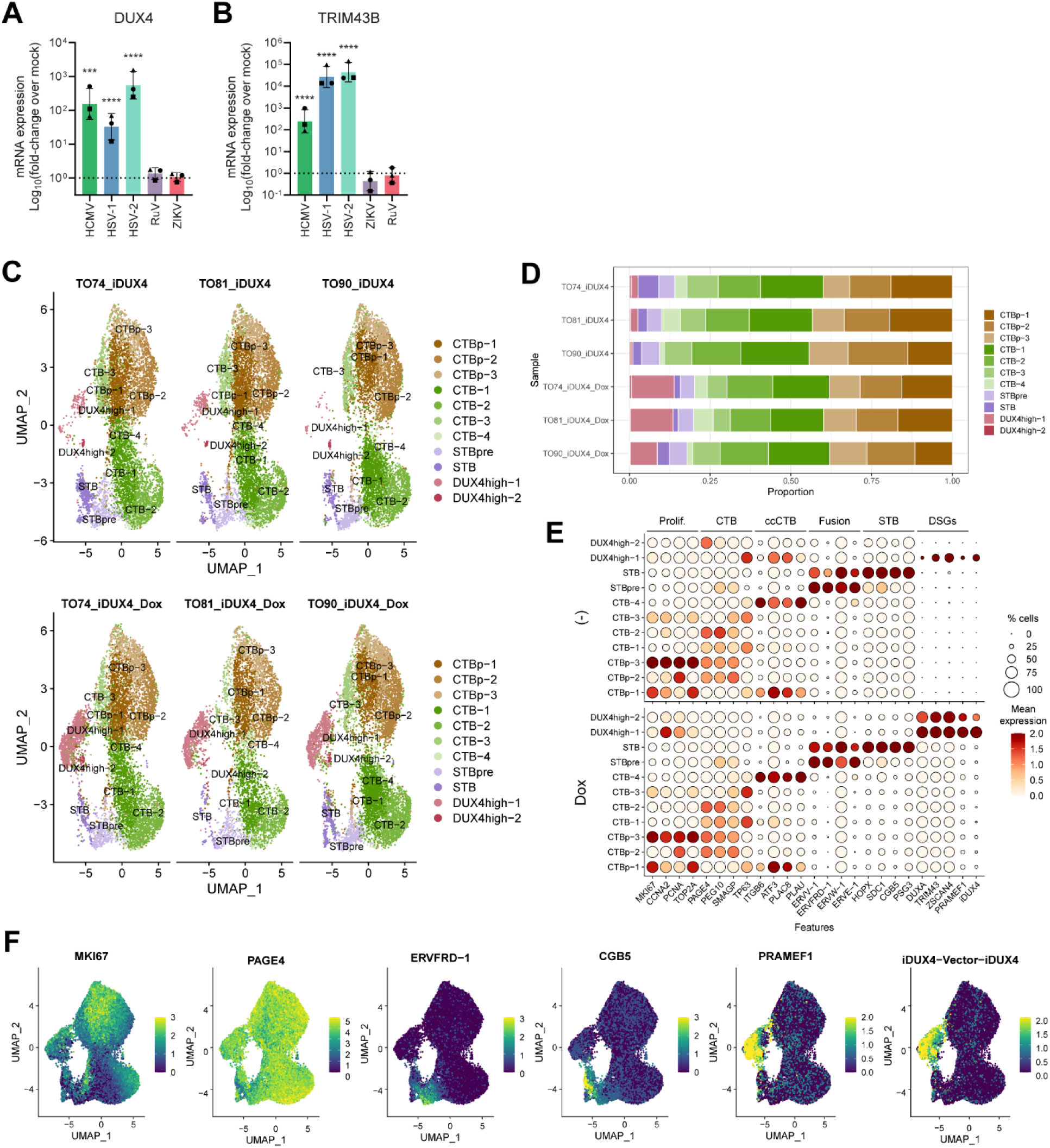
Trophoblasts respond to herpesviruses with DUX4-stimulated genes. **(A-B)** Bar plot portraying the expression of *DUX4* and the DSG *TRIM43* as measured by qRT-PCR following the infection of TOs from three donors with HCMV, HSV-1, HSV-2, RuV, and ZIKV. Values are plotted as Log_10_(fold-change over mock). Shapes denote the donor of each data point. Bars display mean ± SD. Statistical significance was determined by ANOVA with the Tukey multiple comparisons tests. *** p < 0.001, **** p < 0.0001. **(C)** UMAP of clusters from three donors of mock-treated TOs and three donors of Dox-treated TOs separated by sample. Clusters are labeled as proliferative cytotrophoblasts (CTBp), cytotrophoblasts (CTB), pre-syncytiotrophoblasts (STBpre), syncytiotrophoblasts (STB), or DUX4-high cells. Multiple clusters from a single cell type are annotate with a -number after the cluster name. **(D)** Comparison of the proportion of cells that belong to each cluster by sample. **(E)** Dot plots showing the average expression (ALRA assay) and proportion of cells expressing selected genes in each cluster separated into mock-treated samples (top) and Dox-treated samples (bottom). The selected genes are established markers of proliferation, CTBs, cell column CTBs (ccCTBs), fusion, the STB, and DSGs. **(F)** Feature plots of representative markers of each cell type.

**Figure S3.**
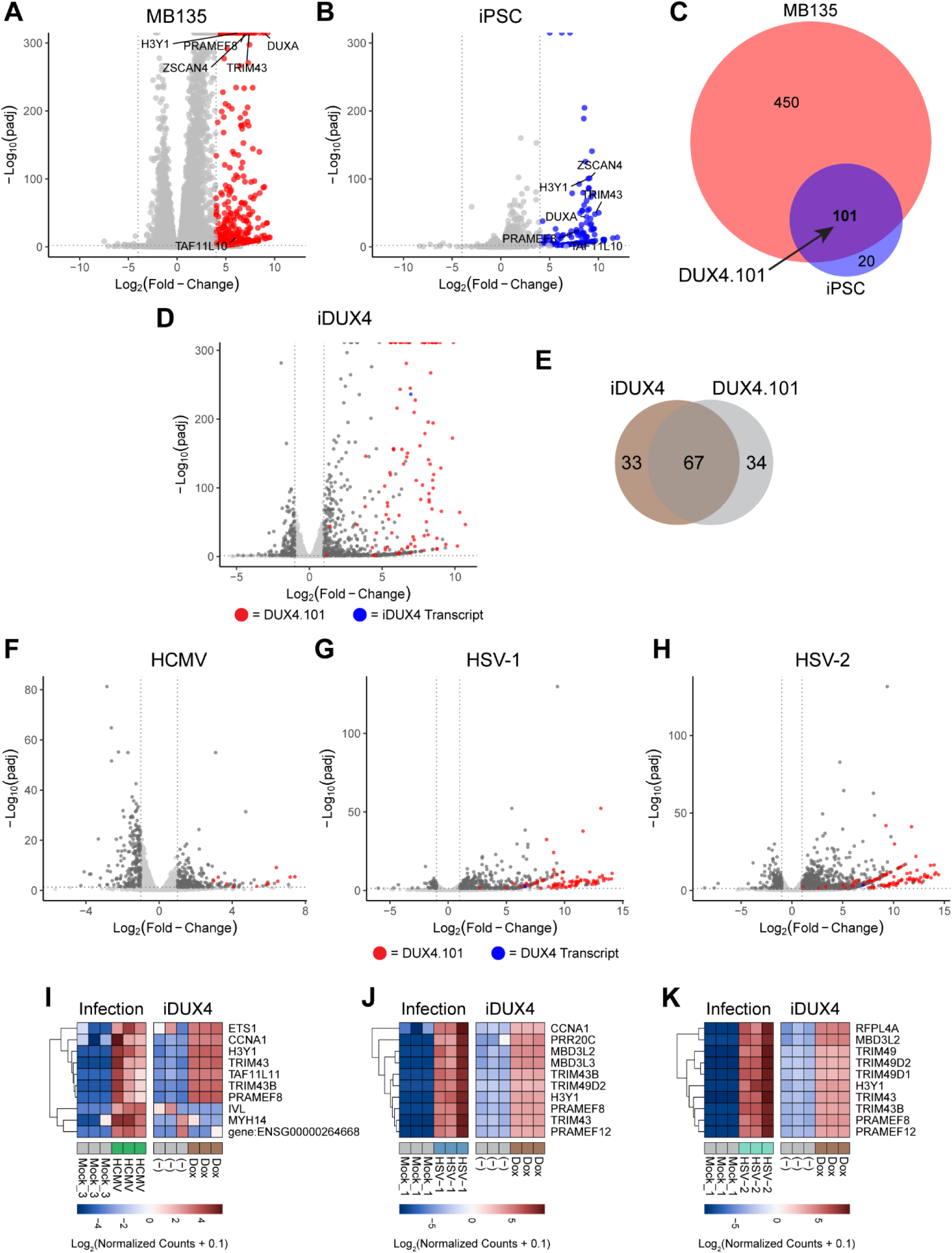
Trophoblasts respond to herpesviruses with DUX4-stimulated genes. **(A-B)** Volcano plots portraying the DEGs upon DUX4 expression in (A) MB135 cells or (B) iPSCs. Colored circles represent significant DEGs (Log2(fold-change) > 4 and adjusted *p*-value < 0.01). Gray circles represent genes whose expression did not meet these criteria. Also see **Table S3**. **(C)** Venn diagram denoting the overlap between the genes induced by DUX4 expression in MB135 cells and iPSCs. These 101 genes constitute the DUX4.101 list used throughout the paper as canonical DSGs that are expressed in divergent cell types. **(D)** TOs engineered to express DUX4 under a doxycycline (Dox) inducible promoter. RNAseq was performed on mock-treated and Dox-treated samples. Volcano plot portrays the DEGs upon DUX4 expression in TOs. Genes that belong to the DUX4.101 list are displayed as red circles and iDUX4 transcript is displayed as a blue circle. Significant DEGs (Log2(fold-change) > 1 and adjusted *p*-value < 0.05) are depicted in dark gray. Light gray was used to depict genes that did not meet these criteria. Also see **Table S3**. **(E)** Venn diagram denoting the overlap between the top 100 DEGs upregulated by iDUX4 expression in TOs and the DUX4.101 list of DSGs. **(F-H)** Volcano plots portraying the DEGs upon HCMV (F), HSV-1 (G), or HSV-2 (H) infection in TOs from Figure 1. Genes that belong to the DUX4.101 list are displayed as red circles and *DUX4* transcript is displayed as a blue circle. Significant DEGs (Log2(fold-change) > 1 and adjusted p-value < 0.05) are depicted in dark gray. Light gray was used to depict genes that did not meet these criteria. **(I-K)** Heatmaps comparing the expression of the top 10 genes induced by HCMV (I), HSV-1 (J), or HSV-2 (K) infection in TOs to the expression of those same genes following iDUX4 expression in TOs. Values are log_2_ normalized counts from DESeq2 analysis plus a pseudo count of 0.1. Keys are at the bottom of each chart. The scale is not constant between each heatmap. Red indicates higher expression and blue indicates lower expression. Hierarchical clustering of genes is shown to the left.

**Figure S4.**
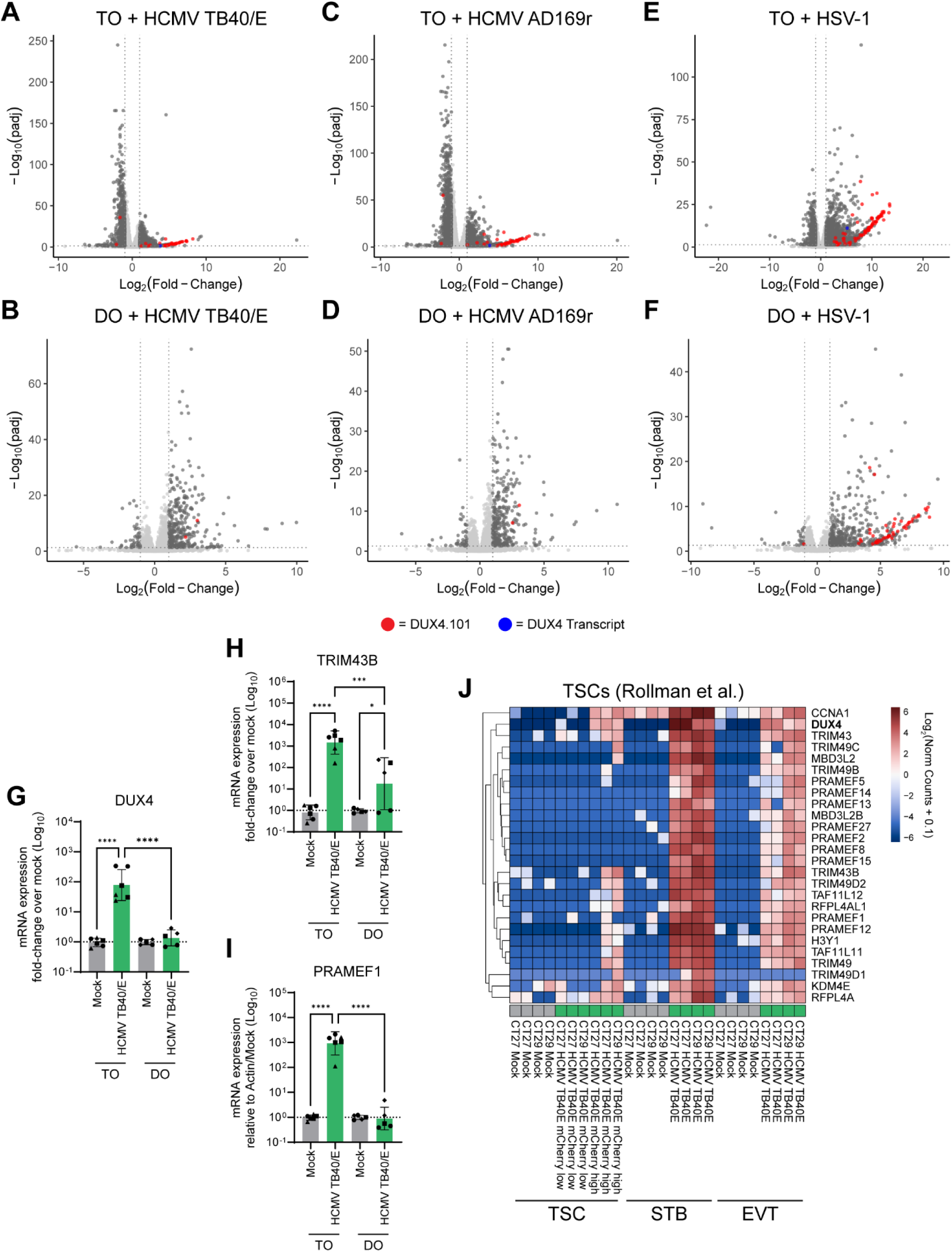
DUX4-stimulated genes are induced to higher levels in trophoblasts than maternal decidual epithelial cells. RNAseq analysis was employed to assess if TOs and Dos differentially express DUX4 or DSGs when infected with HCMV TB40/E, HCMV AD169r, and HSV-1. Also see **Table S5**. **(A-F)** Volcano plots portraying the DEGs upon HCMV TB40/E (A-B), HCMV AD169r (C-D), or HSV-1 (E-F) infection in TOs (top) and DOs (bottom). Genes that belong to the DUX4.101 list are displayed as red circles and DUX4 transcript is displayed as a blue circle. Significant DEGs (Log2(fold-change) > 1 and adjusted p-value < 0.05) are depicted in dark gray. Light gray was used to depict genes that did not meet these criteria. **(G-I)** TOs and DOs from three donors each (two matched and one unique to each) were infected with HCMV TB40/E and qRT-PCR was used to measure the expression of *DUX4* and the DSGs *TRIM43B* and *PRAMEF1*. Values are plotted as Log_10_(fold-change over mock). Shapes denote the donor of each data point. Bars display mean ± SD. Statistical significance was determined by ANOVA with the Tukey multiple comparisons tests. **(J)** RNAseq analysis of mock- and HCMV-infected undifferentiated trophoblast stem cells (TSC), STB-differentiated TSCs, and EVT-differentiated TSCs. Heatmaps portraying the expression of DUX4 and canonical DSGs. Values are log_2_ normalized counts from DESeq2 analysis plus a pseudo count of 0.1. Keys are at the bottom of each chart. Red indicates higher expression and blue indicates lower expression. Hierarchical clustering of genes is shown on the left. * p < 0.05, *** p < 0.001, **** p < 0.0001.

**Figure S5.**
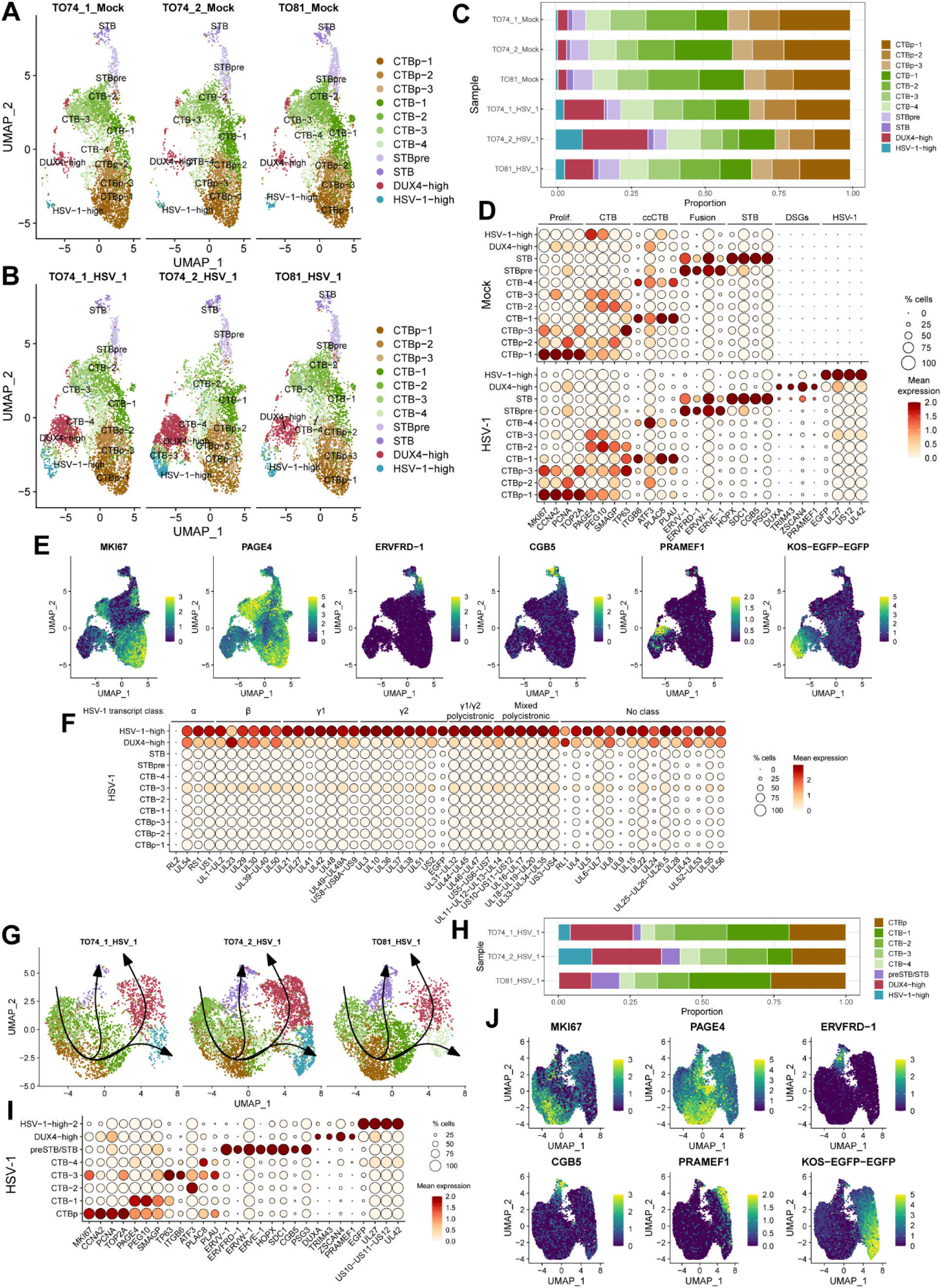
DUX4-stimulated genes define a low viral gene expression trajectory in HSV-1-infected trophoblast organoids. TOs released from matrigel were infected with HSV-1 in suspension then returned to matrigel imbedding. HSV-1-infected and mock-infected organoids from two donors (2x TO74 samples and 1x TO84 sample each) were analyzed by scRNAseq. **(A-B)** UMAP of clusters from all mock-infected (A) and all HSV-1-infected (B) TOs separated by sample. Clusters are labeled as proliferative cytotrophoblasts (CTBp), cytotrophoblasts (CTB), pre-syncytiotrophoblasts (STBpre), syncytiotrophoblasts (STB), DUX4-high cells, or HSV-1-high cells. Multiple clusters from a single cell type are annotate with a -number after the cluster name. **(C)** Comparison of the proportion of cells that belong to each cluster by sample. **(D)** Dot plots showing the average expression (ALRA assay) and proportion of cells expressing selected genes in each cluster separated into mock-infected samples (top) and HSV-1-infected samples (bottom). The selected genes are established markers of proliferation, CTBs, cell column CTBs (ccCTBs), fusion, the STB, DSGs, and HSV-1 transcripts. **(E)** Feature plots of representative markers of each cell type. **(F)** Dot plots showing the average expression (ALRA assay) and proportion of cells expressing all HSV-1 transcripts in each cluster. HSV-1 transcripts are organized into α (immediate early), β (early), γ1 (leaky late), γ2 (true late), γ1/γ2 polycistronic/alternatively spliced, mixed polycistronic/alternatively, and no class. **(G)** UMAP of re-clustered cells from HSV-1 infected samples with >0.1% HSV-1 reads separated by sample. Clusters are labeled as CTBp, CTB, preSTB/STB, DUX4-high, or HSV-1-high. Multiple clusters from a single cell type are annotate with a -number after the cluster name. Slingshot trajectories are superimposed. **(H)** Comparison of the proportion of cells that belong to each cluster by sample. **(I)** Dot plots showing the average expression (ALRA assay) and proportion of cells expressing selected genes in each cluster. The selected genes are established markers of proliferation, CTBs, cell column CTBs (ccCTBs), fusion, the STB, DSGs, and HSV-1 transcripts. **(J)** Feature plots of representative markers of each cell type.

**Figure S6.**
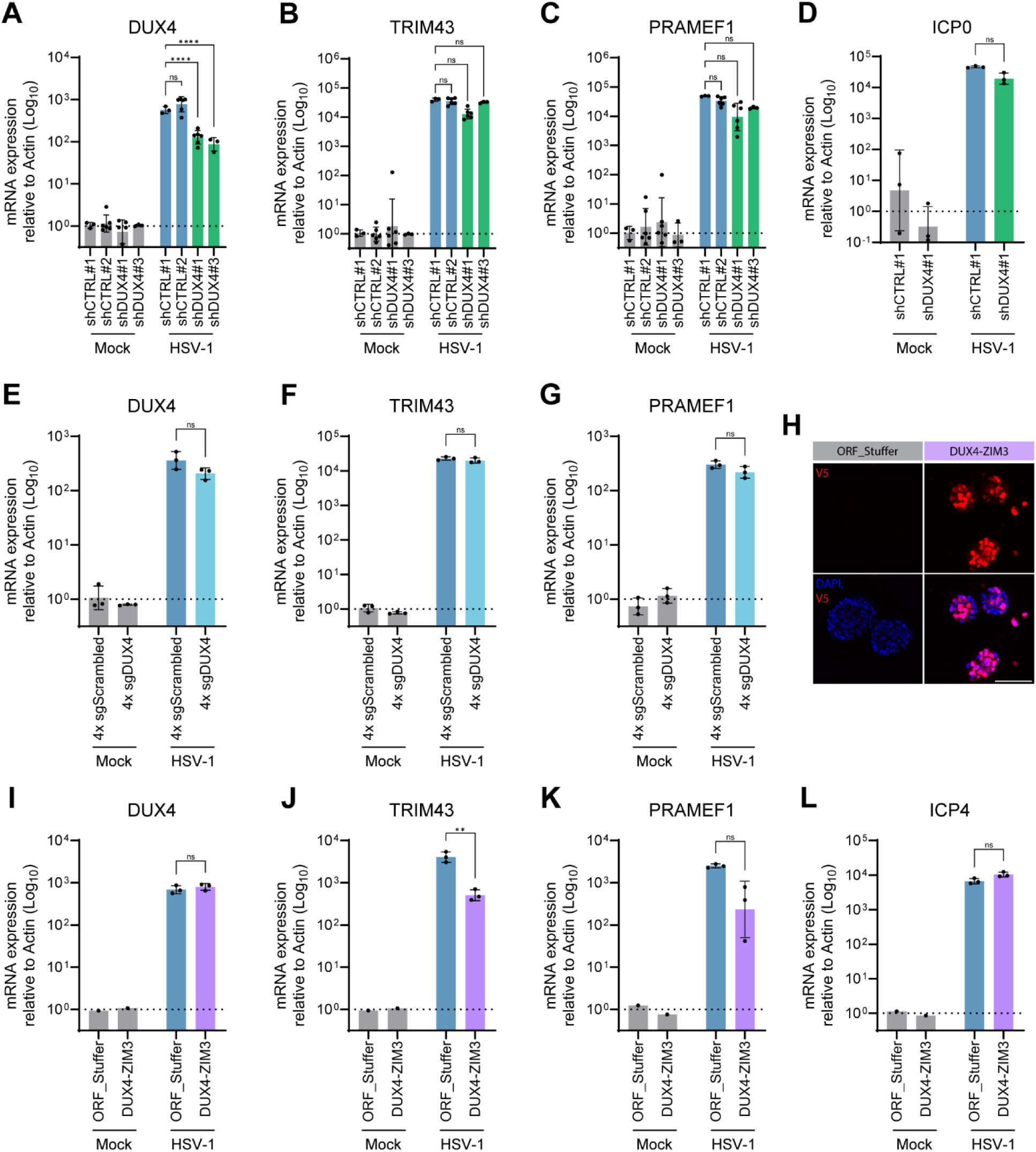
DUX4 knockdown strategies in TOs. **(A-D)** TOs were engineered to express control (shCTRL) or *DUX4*-targeting (shDUX4) shRNAs. Engineered TOs grown on matrigel coats were infected with HSV-1 at an MOI of 5, and RNA was collected at 24 hpi. qRT-PCR was used to measure the expression of *DUX4* (A), *TRIM43* (B), *PRAMEF1* (C), and HSV-1 ICP4 (D). **(E-G)** TOs were engineered to expresses dCas9-KRAB-MeCP2 (used for CRISPRi) along with either 4 scrambled sgRNAs (4x sgScrambled) or 4 sgRNAs targeting the transcription start site of DUX4 (4x sgDUX4). Engineered TOs grown on matrigel coats were infected with HSV-1 at an MOI of 5, and RNA was collected at 24 hpi. qRT-PCR was used to measure the expression of *DUX4* (E), *TRIM43* (F), and *PRAMEF1* (G). **(H-L)** TOs were engineered to express a control vector (ORF_Stuffer) or a dominant-negative DUX4 construct with the transcriptional activation domain of DUX4 replaced with the KRAB domain of ZIM3 along with a V5 tag (DUX4-ZIM3). Confocal micrographs. V5 staining is in red and DAPI-stained nuclei are in blue. Scale bar = 100 μm (H). Engineered TOs grown on matrigel coats were infected with HSV-1 at an MOI of 5, and RNA was collected at 24 hpi. qRT-PCR was used to measure the expression of *DUX4* (I), *TRIM43* (J), *PRAMEF1* (K), and HSV-1 ICP0 (L).

**Figure S7.**
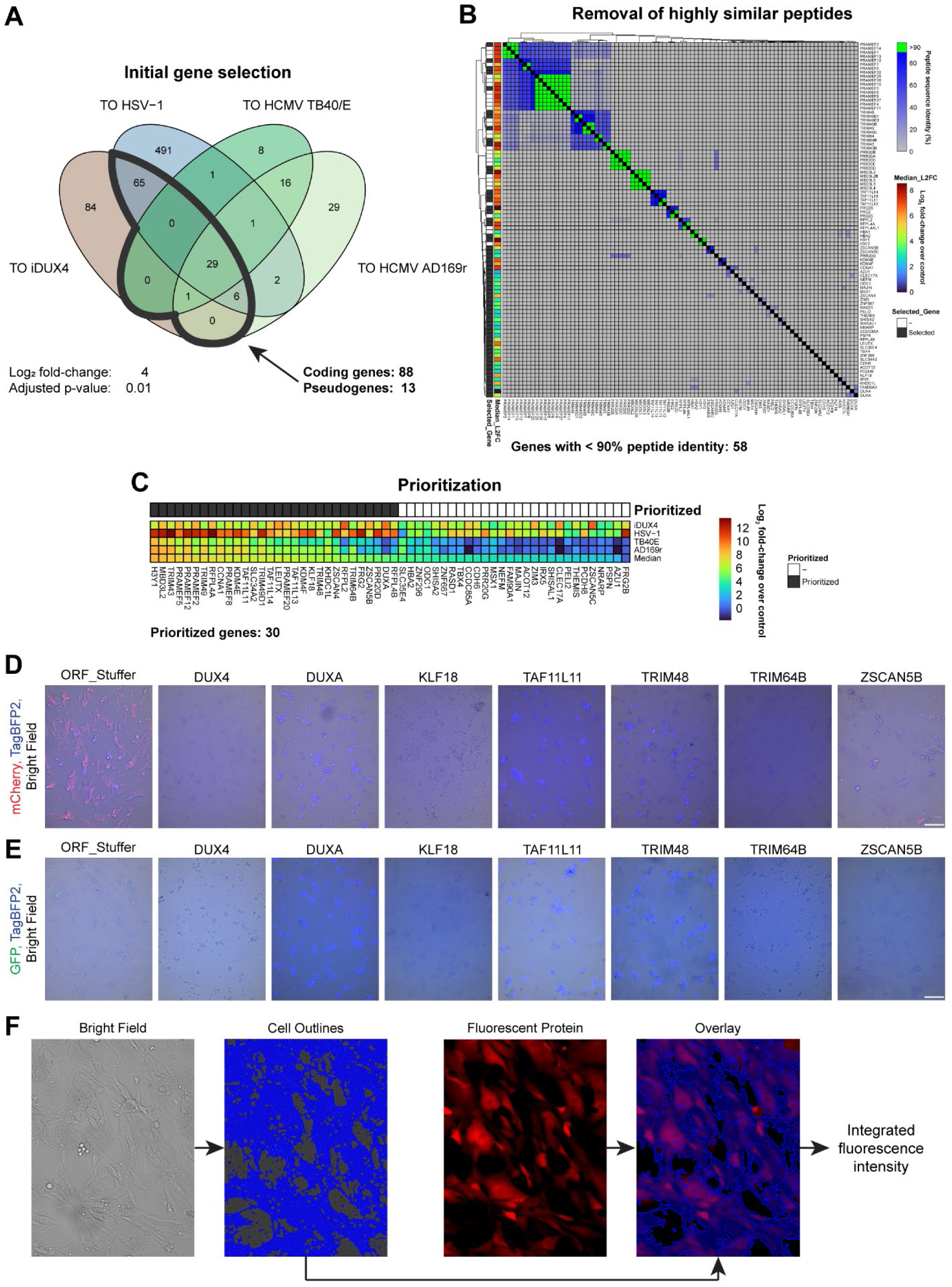
DUX4-stimulated genes exert anti-herpesvirus activity. **(A)** Venn diagram depicting the overlap between DEGs induced by iDUX4 expression or HSV-1, HCMV TB40/E, or HCMV AD169r infection in TOs (Log_2_(fold-change) > 4 and adjusted p-value < 0.01). Thick line highlights the selected genes. **(B)** Heatmap depicting the percent peptide sequence identity between genes selected in (A). Genes with >90% peptide sequence identity are highlighted in green. Median log_2_ fold-change over control from iDUX4 expression and HSV-1, HCMV TB40/E, and HCMV AD169r infection in TOs are displayed to the left. Genes selected for further consideration are marked in gray to the left. **(C)** Heatmap depicting the log_2_ fold-change over control from iDUX4 expression and HSV-1, HCMV TB40/E, and HCMV AD169r infection in TOs. Genes are ordered by median log_2_ fold-change over control. Prioritized genes are marked in gray to the top. **(D-E)** DUX4 and six DSGs were toxic to MRC5 cells. Widefield micrographs of representative fields from HCMV (D) and HSV-1 (E) infection at 24 hpi. Images portray merged TagBFP2 (blue), brightfield (grayscale), and mCherry (D; red; HCMV) or GFP (E; green; HSV-1). **(F)** Schematic of image quantification method. Scale bars = 200 μm.

**Figure S8.**
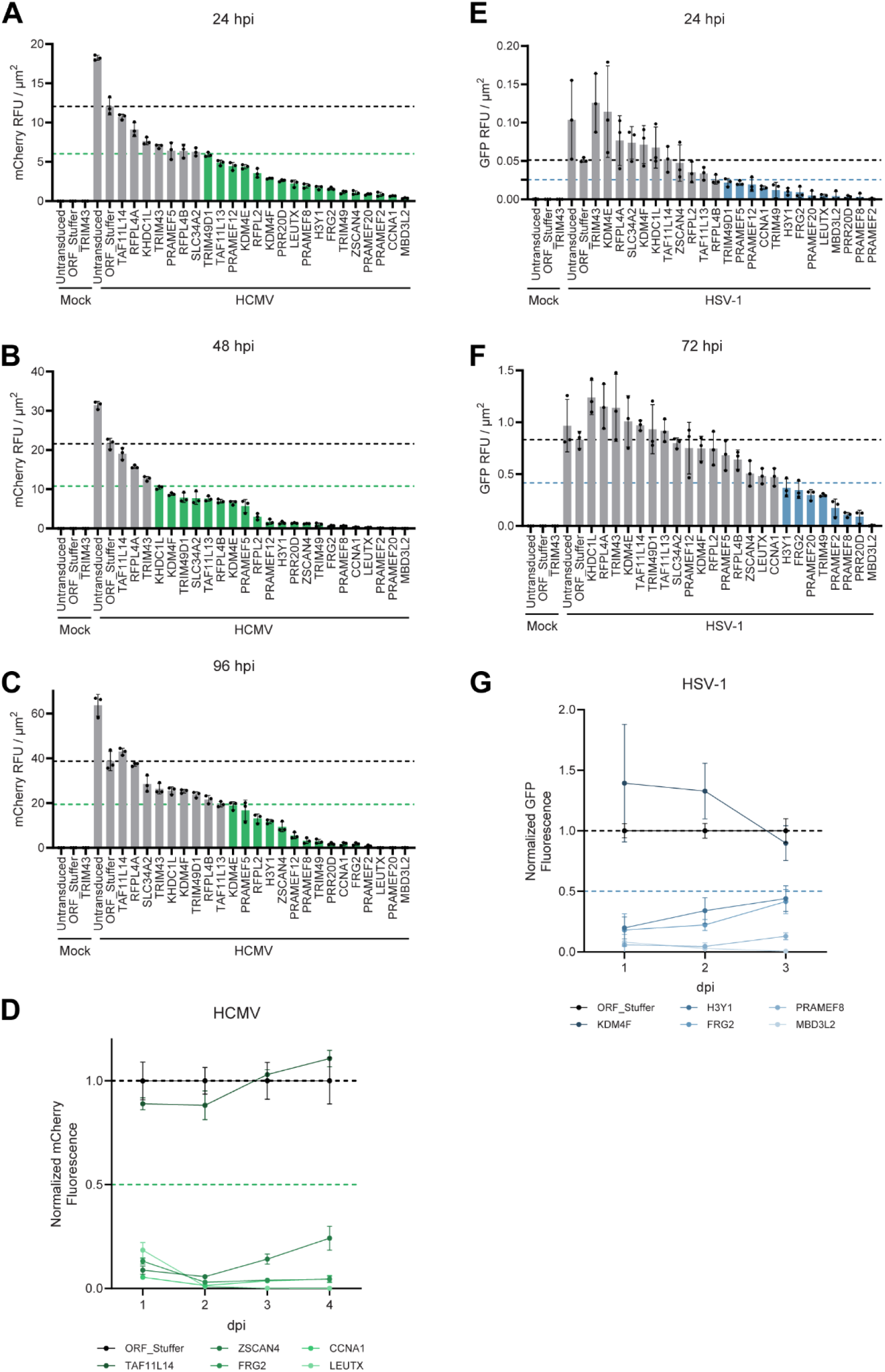
DUX4-stimulated genes exert anti-herpesvirus activity. **(A-C and E-F)** Bar graphs depicting timepoints for HCMV (A-C) and HSV-1 (E-F) not shown in Figure 7. Black circles represent each individual datapoint. Bars display mean ± SD. **(D and G)** Representative DSGs for HCMV: one with minimal impact and four top hits. Normalized mCherry fluorescence from HCMV for these genes at all timepoints (D). Representative DSGs for HSV-1: one with minimal impact and four top hits. Normalized GFP fluorescence from HSV-1 for these genes at all timepoints (G). Dots and error bars display mean ± SD. Black dashed lines in graphs represent the mean expression of ORF_Stuffer. Colored dashed lines represent 0.5x the black line.

